# Endosomal hitchhiking and NDR kinase signaling coordinate SsdA-mRNP localization

**DOI:** 10.64898/2026.04.14.718127

**Authors:** Domenico Modaffari, Edward W.J. Wallace, Kenneth E. Sawin

## Abstract

Transport and regulation of messenger ribonucleoprotein complexes (mRNPs) is critical for spatial control of gene expression within cells. How mRNPs are trafficked in hyphae of different filamentous fungi remains poorly understood. Here we investigate the transport of SsdA, the *Aspergillus nidulans* ortholog of budding yeast RNA-binding protein Ssd1, which is thought to function in translational repression. SsdA forms cytoplasmic puncta that colocalize with the poly(A)-binding protein FabM, marking them as mRNPs, and punctum formation depends on conserved RNA-binding residues of SsdA. SsdA puncta move bidirectionally along microtubules by hitchhiking on early endosomes, and this requires the adaptor proteins PxdA and DipA—machinery previously associated exclusively with peroxisome transport. SsdA puncta are largely depleted from hyphal tip-proximal regions, despite an abundance of early endosomes near hyphal tips, and the length of SsdA puncta depletion zones correlates with hyphal growth rate. We show that acute inhibition of Nuclear Dbf2-related (NDR)-family protein kinase CotA causes rapid accumulation of SsdA puncta near hyphal tips, indicating that puncta depletion depends on CotA activity. Mutation of predicted CotA phosphorylation sites within SsdA to nonphosphorylatable residues also leads to accumulation of SsdA puncta near tips, while mutation to phosphomimetic residues disrupts puncta entirely. Together, our findings establish SsdA-containing mRNPs as a new endosomal hitchhiking cargo in *A. nidulans* and reveal that NDR kinase signaling spatially regulates SsdA-mRNP distribution during normal polarized growth.

## Introduction

Transport of messenger ribonucleoprotein complexes (mRNPs) is a fundamental strategy for spatial control of gene expression in eukaryotic cells (Holt and Bullock, 2009; Martin and Ephrussi, 2009; Bourke et al., 2023). By localizing mRNAs to specific subcellular regions, cells can restrict synthesis of specific proteins to where and when they are needed, supporting processes ranging from embryonic axis specification to synaptic plasticity (Holt and Bullock, 2009; Martin and Ephrussi, 2009).

Cells with compact geometries may be able to distribute mRNAs and their protein products by passive diffusion. However, cells with extended morphologies, such as neurons in animal cells or hyphae in filamentous fungi, face fundamentally different spatial challenges that can require directed transport of these components (Egan et al., 2012a; Sahoo et al., 2018). Microtubule-based transport provides a means for moving specific cargoes over long intracellular distances, in multiple ways (Hirokawa et al., 2009; Reck-Peterson et al., 2018). One important mechanism of transport on microtubules involves motile membrane-bound organelles serving as platforms for the binding and transport of diverse additional cargoes; this process, termed hitchhiking, has been observed in both animal and fungal cells (Baumann et al., 2012; Guimaraes et al., 2015; Salogiannis et al., 2016; Christensen and Reck-Peterson, 2022; Schuhmacher et al., 2023). Much of what is known about hitchhiking is derived from studies in filamentous fungi, in which peroxisomes, lipid droplets, endoplasmic reticulum (ER) and mRNPs have been variously shown to hitchhike on early endosomes (Baumann et al., 2012; Guimaraes et al., 2015; Salogiannis et al., 2016; Christensen and Reck-Peterson, 2022).

In the basidiomycete fungus *Ustilago maydis*, mRNPs hitchhike on early endosomes via the RNA-binding protein (RBP) Rrm4 (Becht et al., 2005; Baumann et al., 2012). However, Rrm4 has no homolog in ascomycete fungi, in which mRNP hitchhiking has been little characterized (Müller et al., 2019). In the filamentous ascomycete *Sordaria macrospora,* the RBP GUL1 has been observed to move with poly(A)-binding protein on early endosomes (Stein et al., 2020). Interestingly, GUL1 is the *S. macrospora* ortholog of the well-studied *Saccharomyces cerevisiae* RBP Ssd1, which is thought to function as a translational repressor (Uesono et al., 1997; Jansen et al., 2009; Ballou et al., 2021; Bayne et al., 2022).

Here, using the model filamentous ascomycete *Aspergillus nidulans*, we show that SsdA, the *A. nidulans* ortholog of Ssd1/GUL1, hitchhikes on early endosomes, and we uncover mechanistic features of hitchhiking and of SsdA-mRNP regulation in relation to hyphal tip growth. SsdA forms cytoplasmic puncta that move bidirectionally along microtubules by hitchhiking on early endosomes. Further evidence indicates that SsdA puncta represent mRNPs, and their transport requires adaptor proteins PxdA and DipA, previously implicated in peroxisome hitchhiking on early endosomes (Salogiannis et al., 2016, 2021). We further demonstrate that SsdA-mRNP puncta are depleted in tip-proximal hyphal regions, and the extent of depletion is correlated with hyphal growth rate. Using an analog-sensitive kinase approach and SsdA phosphorylation-site mutants, we show that tip-proximal depletion of SsdA-mRNP puncta depends on the activity of the Nuclear Dbf2-related (NDR) kinase CotA (the *A. nidulans* ortholog of *S*. *cerevisiae* Cbk1) near the hyphal tip, acting on NDR kinase consensus sites within the N-terminus of SsdA. Collectively, our findings on transport and regulation of SsdA-mRNP puncta reveal that different endosomal hitchhiking cargoes may share a common machinery for hitchhiking, and that NDR kinase activity, which has been previously reported to regulate sequestration of mRNPs into storage granules in association with cell stress (Jansen et al., 2009; Kurischko et al., 2011a; Nuñez et al., 2016), can also regulate mRNP spatial distribution during normal hyphal growth.

## Results

### SsdA is the *A. nidulans* homolog of *S. cerevisiae* RNA-binding protein Ssd1 and forms motile cytoplasmic puncta

The *A. nidulans* gene AN1158 was previously identified as a homolog of *S. cerevisiae* Ssd1 (Jansen et al., 2009). Following *Aspergillus* nomenclature, we named AN1158 as *ssdA*. AlphaFold3 (Abramson et al., 2024) predicts SsdA’s structured domains with high confidence (**Fig. S1A**), and the predicted SsdA structure closely resembles the Ssd1 crystal structure (**Fig. 1A–C**), in line with Ssd1 sequence conservation across Pezizomycotina (Ballou et al., 2021). The only notable differences between the two structures were associated with two insertions in SsdA relative to Ssd1 (**Fig. 1A–C**).

**Figure 1.**
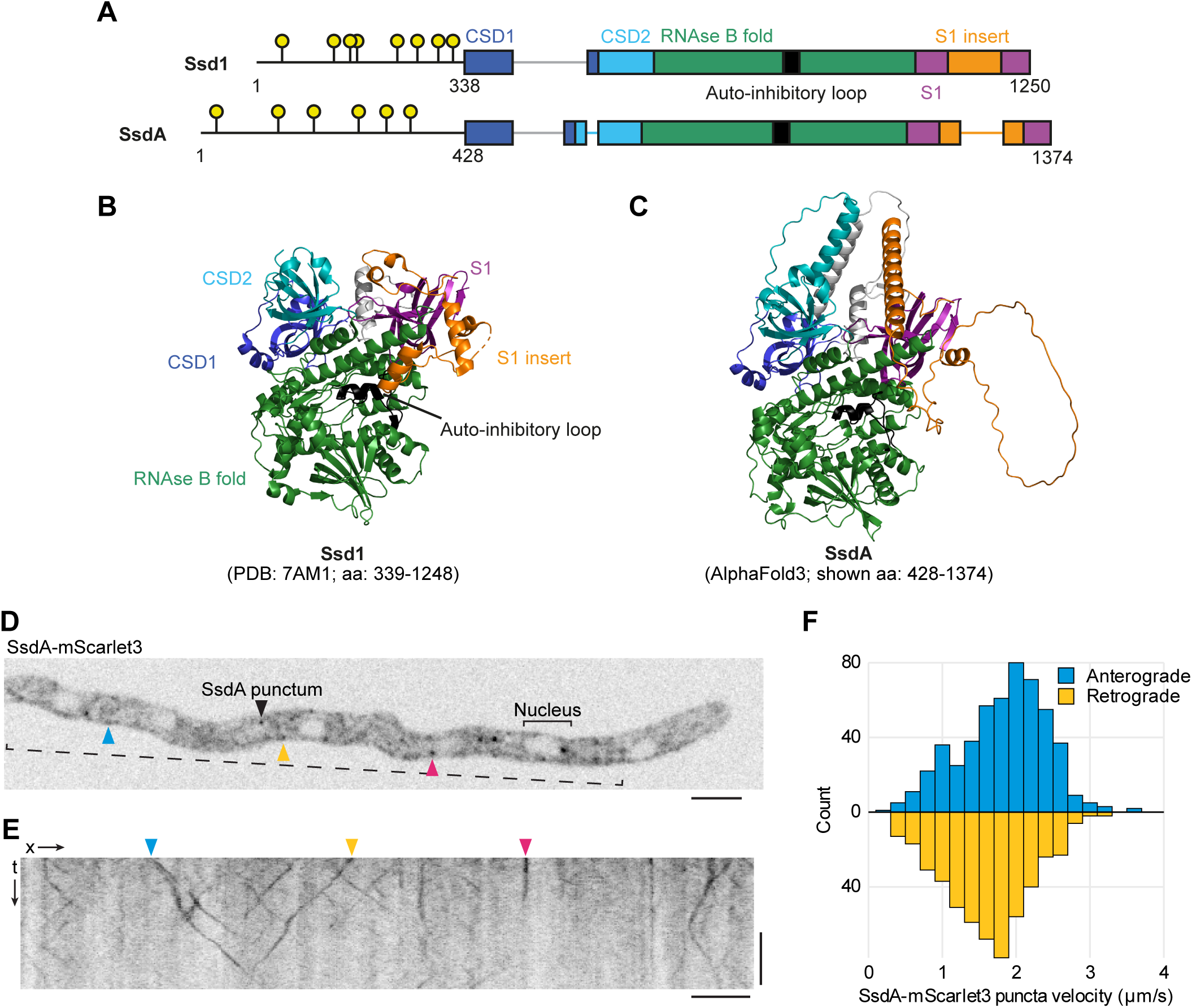
SsdA, the *Aspergillus nidulans* ortholog of budding yeast RNA-binding protein Ssd1, localizes to motile puncta. **(A)** Domain maps of *S. cerevisiae* Ssd1 and *A. nidulans* SsdA. Circles indicate predicted Cbk1/NDR kinase phosphorylation sites (Jansen et al., 2009). *A. nidulans* phosphorylation sites were identified according to the Cbk1/NDR consensus motif H-X- (K/R)-(K/R)-X-(S/T) (Mazanka et al., 2008). **(B)** Structure of *S. cerevisiae* Ssd1, color-coded as in Bayne et al. (2022). An ∼70 amino-acid region within the CSD1 domain was not resolved and is absent from the structure. **(C)** AlphaFold3 model of *A. nidulans* SsdA, color-coded by domain as in (B). Regions of CSD2 and S1 insert domains not present in Ssd1 were modelled with lower confidence (see Fig. S1A). **(D, E)** Zero timepoint from single-z-section continuous-acquisition movie of hypha expressing SsdA-mScarlet3 from endogenous locus (D), with accompanying kymograph (E). SsdA-mScarlet3 signal is present in the cytosol in both punctate and diffuse form but is absent from nuclei. Bracket indicates kymographed region shown in (E). Arrowheads indicate examples of puncta colored by movement state: blue, anterograde (towards tip); yellow, retrograde (away from tip); magenta, immotile. **(F)** Velocity distribution of SsdA-mScarlet3 puncta. Velocities were derived from 10 s kymographs using Kymobutler (Jakobs et al., 2019). Pooled data from 3 biological replicates; number of movements scored = 1025. Colors indicate direction of movement. Scale bars = 5 µm, 2 s.

We first asked whether Ssd1’s mRNA-binding specificity may be conserved in *A. nidulans*. To test this, we searched 5′ untranslated region (UTR) sequences from the *A. nidulans* transcriptome for the Ssd1 consensus binding motif “CNYUCNYU”, which is enriched in 5′ UTRs of Ssd1 target mRNAs (Bayne et al., 2022). We found that this motif is present in 5′ UTRs of over 2000 *A. nidulans* mRNAs (**Fig. S1B**), and it is particularly enriched in 5′ UTRs of mRNAs encoding cell wall and cell polarity proteins (**Fig. S1C**). This distribution is consistent with *A. nidulans* SsdA having target mRNA specificity similar to that of *S. cerevisiae* Ssd1 (see Discussion).

To investigate SsdA localization *in vivo*, we used a strain expressing SsdA-mScarlet3 from the endogenous locus (Modaffari et al., 2024). Tagging *ssdA* with mScarlet3 did not affect colony morphology, indicating that the tagged protein is likely functional (**Fig. S1D**). By contrast, deletion of *ssdA* produced smaller colonies with a sharp edge (**Fig. S1D**). Live-cell imaging revealed that SsdA-mScarlet3 was present in a diffuse cytosolic pool and as discrete motile puncta (**Fig. 1D, E; Video S1)**. SsdA-mScarlet3 was largely excluded from nuclei, although stationary puncta were sometimes observed at juxtanuclear positions.

SsdA puncta exhibited bidirectional movement along the hyphal axis, with a mean velocity of ∼1.8 µm/s (**Fig. 1E, F; Video S1**). Anterograde and retrograde movements occurred at roughly equal frequency (**Fig. 1F**). Puncta frequently moved out of the focal plane during single z-plane imaging, preventing accurate assessment of processivity. To confirm that puncta formation was not an artifact of the mScarlet3 tag, we also tagged *ssdA* with the green fluorescent protein mNeonGreen. SsdA-mNeonGreen also formed motile puncta and a diffuse cytoplasmic pool (**Fig. S1E, F; Video S1**), demonstrating that this localization pattern represents *bona fide* SsdA behavior rather than a tag-specific artifact.

### Moving SsdA puncta represent messenger ribonucleoprotein complexes

Because SsdA is likely an RBP, we hypothesized that SsdA puncta represent mRNP complexes. In *S. cerevisiae*, Ssd1 interacts with the poly(A)-binding protein Pab1 (Richardson et al., 2012), a core mRNP component and widely used cytological marker for mRNA (Hogan et al., 2008; König et al., 2009). In *S. macrospora,* the SsdA ortholog GUL1, which shows a similar localization pattern to SsdA, also colocalizes with Pab1 (Stein et al., 2020). We therefore asked whether SsdA associates with FabM, the *A. nidulans* Pab1 ortholog.

FabM-mScarlet3, tagged at the endogenous locus, appeared predominantly cytosolic (**Fig. S2A; Video S2**), consistent with previous reports (Soukup et al., 2017). This made it difficult to determine whether SsdA puncta colocalize with FabM. To reveal any motile punctate pool of FabM that might be obscured by high cytosolic signal, we used fluorescence recovery after photobleaching (FRAP). Interestingly, after photobleaching of FabM within defined regions of hyphae, we observed both anterograde and retrograde movements of FabM puncta passing through the bleached region (**Fig. S2A–C; Video S2**).

To determine whether SsdA localizes to FabM-containing mRNPs, we used the same FRAP approach to image a strain coexpressing SsdA-mNeonGreen and FabM-mScarlet3. Kymographs of photobleached regions revealed that SsdA and FabM puncta moved at indistinguishable velocities, consistent with their co-transport (**Fig. S2D**). In addition, FabM and SsdA puncta movements strongly colocalized (**Fig. 2A, B**); about 90% of detectable FabM puncta contained SsdA (**Fig. S2E**), and there was negligible spectral bleed-through between channels (**Fig. S2C**). Because we cannot be confident of FabM puncta detection efficiency after photobleaching, we did not quantify the fraction of SsdA puncta that contain FabM.

**Figure 2.**
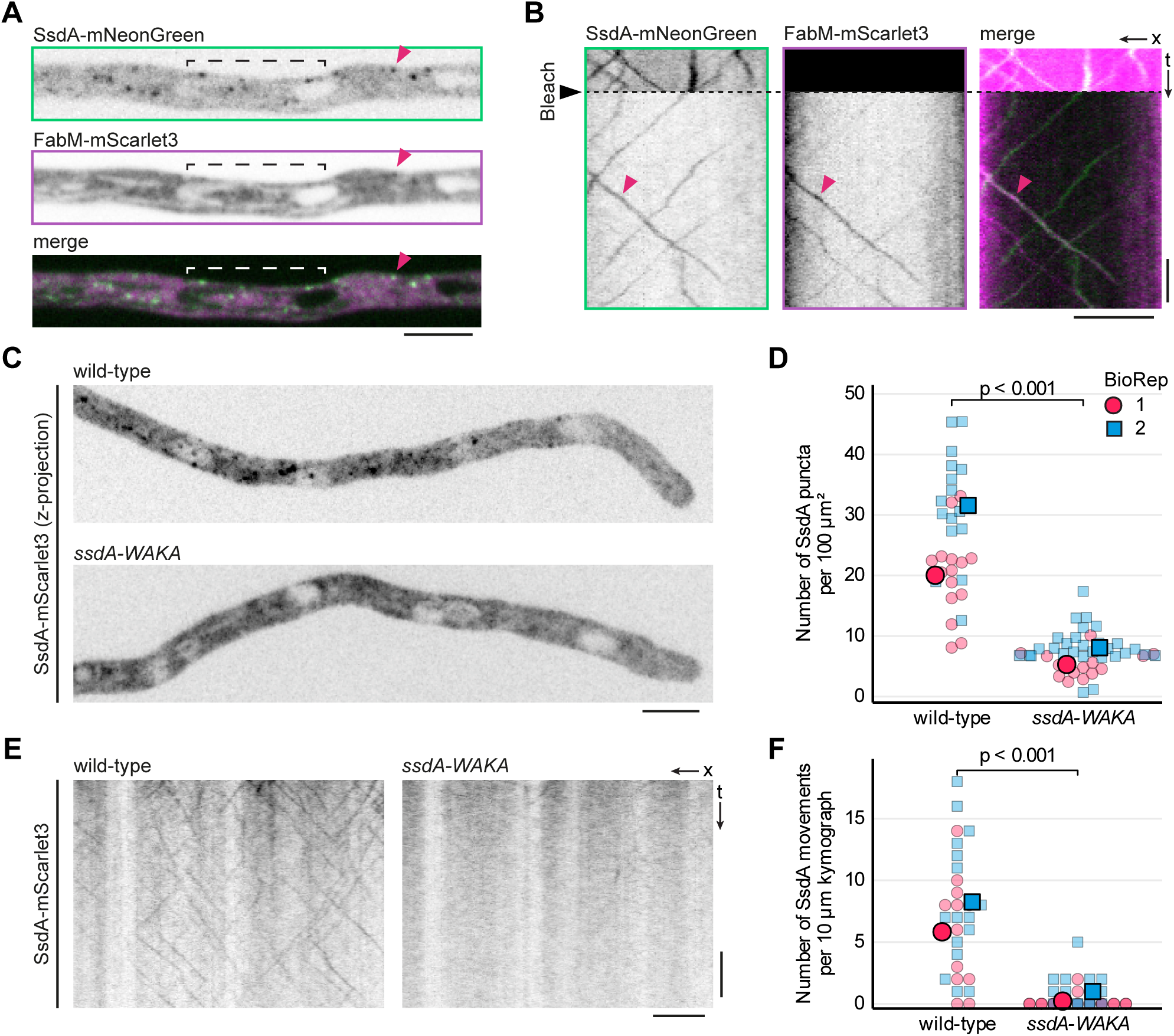
SsdA puncta colocalize with mRNP marker poly(A)-binding protein FabM, dependent on conserved SsdA residues implicated in RNA binding. **(A)** Zero timepoint from single-z-section simultaneous dual-channel movie of hypha expressing SsdA-mNeonGreen and poly(A)-binding protein FabM-mScarlet3, prior to photobleaching. Brackets indicate prospective bleached region related to kymograph in (B). Arrowheads indicate colocalizing puncta. **(B)** Kymographs from the movie associated with (A), spanning the bracketed bleach region. Magenta arrowheads indicate colocalizing puncta movements. **(C)** Maximum-intensity z-projection micrographs of hyphae expressing wild-type and W652A, K654A (WAKA) SsdA-mScarlet3. **(D)** SsdA puncta area number density in hyphae expressing wild-type or WAKA mutant SsdA-mScarlet3. Small symbols indicate individual hyphae; large symbols indicate means within biological replicates (BioReps). Different shapes/colors indicate distinct BioReps. Significance brackets indicate thresholds met in both BioReps (Mann–Whitney *U*, two-tailed, per BioRep). Number of hyphae: wild-type = 15, 16; *ssdA-WAKA* = 14, 30 for BioRep 1, 2 respectively. **(E)** Representative kymographs from single-z-section continuous-acquisition movies of wild-type and WAKA mutant SsdA-mScarlet3 strains. **(F)** Number of SsdA-mScarlet3 movements per kymograph in hyphae expressing wild-type or WAKA mutant SsdA-mScarlet3. Each kymograph (10 s duration) was derived from a 10 µm line adjacent to the third most tip-proximal nucleus. Small symbols indicate individual kymographs; large symbols indicate means within BioReps. Different shapes/colors indicate distinct BioReps. Significance brackets indicate thresholds met in both BioReps (Mann–Whitney *U*, two-tailed, per BioRep). Number of kymographs: wild-type = 12, 16; *ssdA-WAKA* = 18, 16 for BioRep 1, 2 respectively. Data for wild-type are copied in Figure 4B, as data for wild-type and all mutants were collected in the same experiments (see Methods). Scale bars = 5 µm, 2 s.

To test whether RNA binding is required for SsdA localization to puncta, we mutated SsdA residues predicted to be important for RNA binding. In *S. cerevisiae* Ssd1, the double mutant W583A, K585A (“CSD-TOP” mutant) decreases RNA binding ∼33-fold (Bayne et al., 2022). We therefore mutated the comparable residues (W652A and K654A) in the *ssdA::mScarlet3* gene, we refer to this mutant as *ssdA-WAKA* (**Fig. S2F**). The *ssdA-WAKA* mutation phenocopied *ssdAΔ* colony morphology, consistent with loss of function (**Fig. S2G**). This phenotype was not due to decreased expression, as SsdA-WAKA protein levels were similar to wild type (**Fig. S2H, I**). Using a fine-tuned Spotiflow model to ensure consistent, unbiased puncta detection across conditions (Dominguez Mantes et al., 2025; see Methods), live-cell imaging revealed that *ssdA-WAKA* hyphae contained about 3-fold fewer SsdA puncta than wild-type (**Fig. 2C, D**). In kymographs, most detectable puncta movement was lost (**Fig. 2E, F**), although when SsdA-WAKA puncta could be observed, they were motile. We conclude that moving SsdA puncta represent mRNP complexes, and that SsdA localization to moving puncta depends on its RNA-binding activity.

To test the possibility that SsdA might be required for transport of mRNPs more generally, we deleted the *ssdA* gene in a strain expressing FabM-mScarlet3. In *ssdAΔ* hyphae, FabM puncta remained motile, with unchanged velocity relative to wild-type (**Fig. S2J–L**). This indicates that SsdA is dispensable for general mRNP transport. We did not quantify overall number of FabM movements because of the uncertain efficiency of FabM puncta detection by FRAP.

### SsdA puncta hitchhike on early endosomes using hitchhiking adaptors PxdA and DipA

Because SsdA puncta moved at velocities reminiscent of microtubule-based early endosome transport in *A. nidulans* (Abenza et al., 2009; Zekert and Fischer, 2009), we tested whether SsdA puncta move on microtubules. We imaged SsdA-mScarlet3 together with GFP-tagged α-tubulin (TubA-GFP; Horio and Oakley, 2005; Qiu et al., 2022). Time projections showed that SsdA puncta followed microtubule tracks (**Fig. 3A; Fig. S3A; video S3**). To test whether microtubules are required for SsdA puncta motility, we depolymerized microtubules using methyl benzimidazol-2-yl carbamate (MBC).

**Figure 3.**
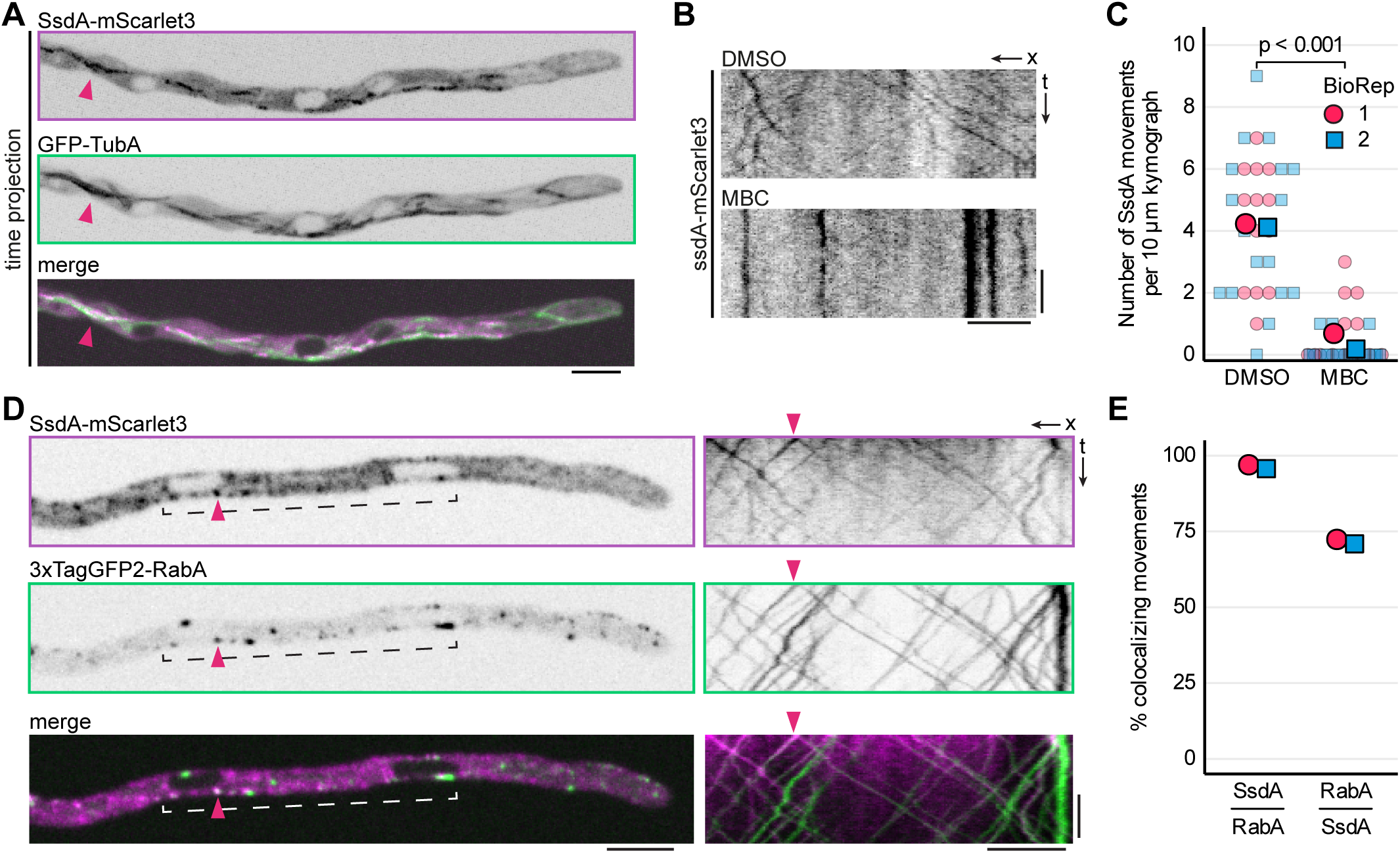
SsdA puncta movement depends on microtubules and is associated with early endosomes. **(A)** Maximum-intensity time projection of hypha expressing both SsdA-mScarlet3 and TubA-GFP. Projection was generated from a 10 s single-z-section continuous-acquisition movie; arrowheads indicate example of colocalization. **(B)** Kymographs of mScarlet3 channel from single-z-section movies of hyphae expressing SsdA-mScarlet3 and TubA-GFP, 10 minutes after exchange to fresh medium containing either 25 µg/mL microtubule-depolymerizing drug MBC or vehicle (DMSO) alone. Micrographs encompassing the kymographed regions and showing both SsdA-mScarlet3 and TubA-GFP channels are shown in Fig. S3B. **(C)** Numbers of SsdA-mScarlet3 movements after MBC or vehicle (DMSO) treatment as in (B). Movements were scored from 10 µm, 5 s kymographs drawn adjacent to a nucleus within each hypha. Small symbols indicate individual kymographs; large symbols indicate means within biological replicates (BioReps). Different shapes/colors indicate distinct BioReps. Significance brackets indicate thresholds met in both BioReps (Mann–Whitney *U*, two-tailed, per BioRep). Sample sizes: vehicle = 13, 17; MBC = 13, 16, for BioRep 1, 2 respectively. **(D)** Zero timepoint from single-z-section simultaneous dual-channel continuous-acquisition movie of hypha expressing SsdA-mScarlet3 and early endosome marker 3xTagGFP2-RabA, with corresponding kymographs at right. Brackets indicate kymographed region. Arrowheads indicate an example of colocalization. **(E)** Percentage of SsdA-mScarlet3 movements colocalizing with 3xTagGFP2-RabA movements (“SsdA/RabA”) and percentage of 3xTagGFP2-RabA movements colocalizing with SsdA-mScarlet3 movements (“RabA/SsdA”), quantified from kymographs as in (D). Number of movements: SsdA = 397, 397; RabA = 532, 533 for BioRep 1, 2 respectively. Scale bars = 5 µm, 2 s.

MBC largely eliminated cytoplasmic microtubules (**Fig. S3B, C**) and led to a strong decrease in SsdA puncta movement (**Fig. 3B, C**). Together, these results show that SsdA puncta undergo microtubule-dependent transport.

In filamentous fungi, early endosomes serve as platforms for long-range microtubule-based transport of mRNPs and some organelles (Baumann et al., 2012; Higuchi et al., 2014; Guimaraes et al., 2015; Salogiannis et al., 2016; Kwon et al., 2020; Christensen and Reck-Peterson, 2022; Vázquez-Carrada et al., 2026). To test whether SsdA puncta associate with early endosomes, we co-imaged SsdA-mScarlet3 with 3xTagGFP2-RabA, the *A. nidulans* Rab5 homolog and a marker of early endosomes (Abenza et al., 2009; Egan et al., 2012b). SsdA and RabA puncta showed considerable colocalization in still images and videos (**Fig. 3D; Video S4**). The only exception to this was in regions very close to hyphal tips (**Fig. S3E**); we address this observation further below.

Velocities of motile SsdA and RabA puncta (i.e. early endosomes) were statistically indistinguishable (**Fig. S3D**), and kymograph analysis revealed extensive colocalization of movement (**Fig. 3D**). Almost all observed SsdA puncta movements colocalized with RabA movements (**Fig. 3E**), indicating that motile SsdA puncta are invariably associated with early endosomes. Conversely, approximately 75% of early endosome movements colocalized with detectable SsdA movements (**Fig. 3E**). Collectively, these results indicate that motile SsdA puncta hitchhike on early endosomes.

Although motile SsdA and RabA puncta colocalized extensively, in a separate analysis of all detectable puncta (see Methods), we found that only approximately one-third of all quantified SsdA puncta colocalized with RabA (**Fig. S3F**). In addition, we noted that SsdA puncta were generally brighter when they colocalized with RabA (**Fig. S3F**). However, when SsdA and RabA did colocalize, only a very weak correlation in their fluorescence signals was observed (**Fig. S3G**). Overall, these results suggest that SsdA can form puncta when not associated with early endosomes, but such puncta may be too faint to be observed readily in kymographs. Moreover, although association with early endosomes may promote SsdA coalescence into puncta, this does not scale directly with RabA abundance.

In *A. nidulans*, peroxisomes hitchhike on early endosomes by associating with endosomal adaptor proteins PxdA and DipA (Salogiannis et al., 2016, 2021). We hypothesized that SsdA puncta might use these same adaptor proteins for hitchhiking. Consistent with this, deletion of either *pxdA* or *dipA* in the SsdA-mScarlet3 strain nearly abolished SsdA puncta movement (**Fig. 4A, B**). This effect was not due to decreased SsdA expression, as western blotting confirmed similar SsdA-mScarlet3 levels across wild-type and deletion strains (**Fig. S4A, B**). Interestingly, deletion of *pxdA* or *dipA* also led to decreased numbers of detectable SsdA puncta, although the decrease was less severe than that observed in the *ssdA-WAKA* mutant (**Fig. S4C**). In addition, particularly bright SsdA puncta, which were readily observed in wild-type hyphae, were largely absent in both *pxdA* and *dipA* mutants (**Fig. S4D**). These observations suggest that *pxdA-* and/or *dipA*-dependent association of SsdA with early endosomes may promote the coalescence or concentration of SsdA-containing mRNPs into larger assemblies.

**Figure 4.**
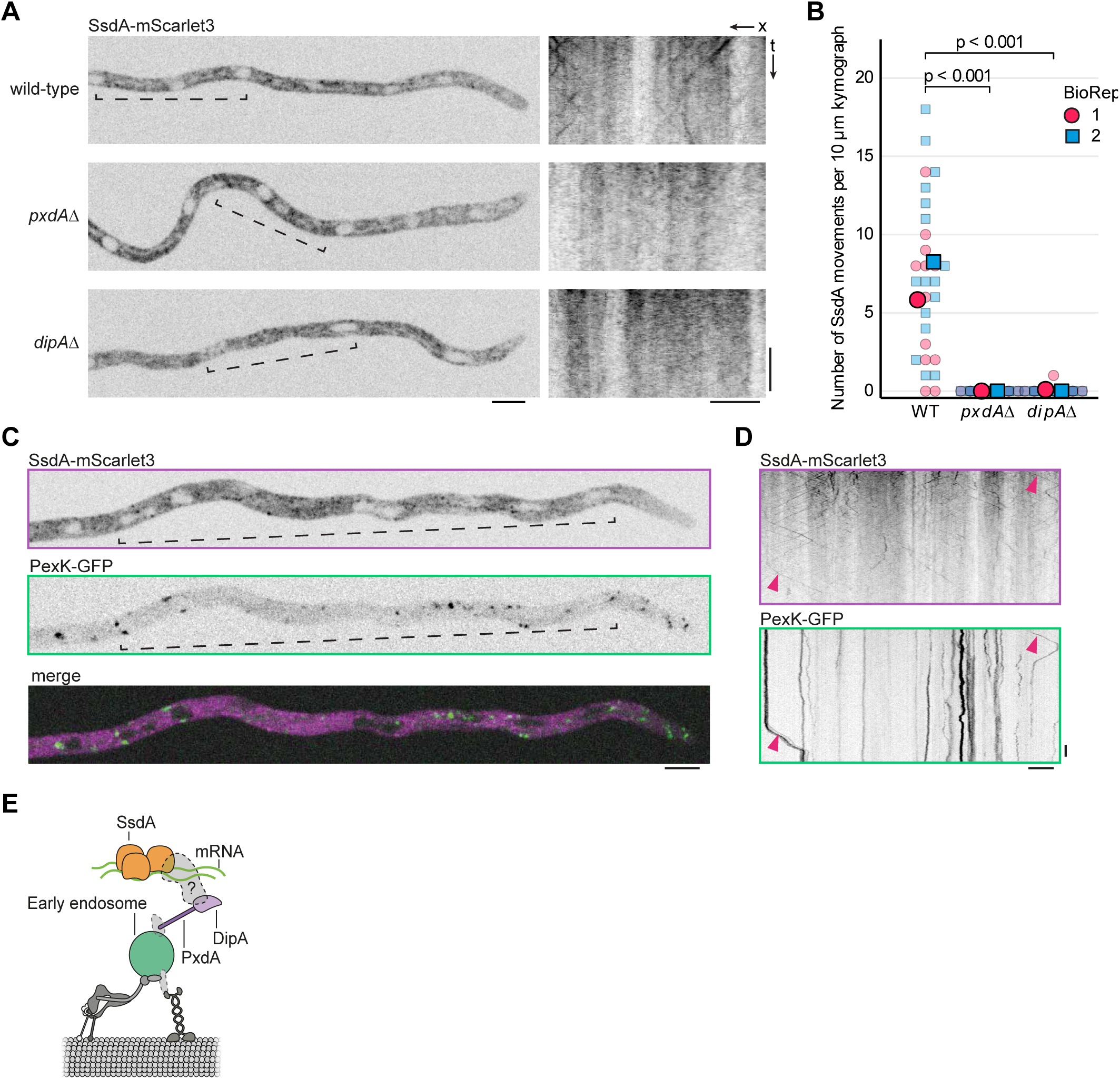
SsdA puncta movement requires peroxisome hitchhiking adaptors PxdA and DipA but is independent of peroxisomes. **(A)** Zero timepoints from single-z-section continuous-acquisition movies of SsdA-mScarlet3 in wild-type and deletion-mutants of hitchhiking adaptors PxdA and DipA, with corresponding kymographs shown at right. Brackets indicate regions used for kymographs. **(B)** Number of SsdA-mScarlet3 puncta movements per kymograph in wild-type, *pxdAΔ*, and *dipAΔ* hyphae. Each kymograph (10 s duration) was derived from a 10 µm line adjacent to the third most tip-proximal nucleus. Data for wild-type are copied from Figure 2F, as data for wild-type and all mutants were collected in the same experiments (see Methods). Significance brackets indicate thresholds met in both biological replicates (BioReps) (Mann–Whitney *U*, two-tailed, per BioRep). Number of kymographs: wild-type = 12, 16; *pxdAΔ* = 13, 12; *dipAΔ* = 10, 15 for BioRep 1, 2 respectively. **(C, D)** Zero timepoint from near-simultaneous single-z-section time-lapse (150 ms interval) movie of hypha expressing SsdA-mScarlet3 and peroxisome marker PexK-GFP (C), with accompanying kymographs (D). Brackets in (C) indicate region kymographed in (D). Arrowheads indicate examples of colocalizing movements. **(E)** Model of SsdA mRNP association with *A. nidulans* endosomal hitchhiking machinery (Salogiannis et al., 2016; Christensen and Reck-Peterson, 2022). Our data indicate that SsdA puncta move with early endosomes and that SsdA puncta movement depends on both PxdA and DipA. Both PxdA and DipA are also required for normal levels of detectable SsdA puncta. Gray shapes represent possible additional bridging factors that may be required for attachment of SsdA-mRNPs and/or PxdA/DipA to early endosomes. Normal levels of SsdA puncta are also compromised in the predicted RNA-binding mutant *ssdA-WAKA*.

PxdA and DipA have previously been implicated exclusively in the transport of peroxisomes and Woronin bodies, which are derived from peroxisomes (Salogiannis et al., 2016, 2021; Songster et al., 2023). This raised the question of whether SsdA puncta are carried as cargoes on peroxisomes or simply use the same adaptor proteins to independently hitchhike on early endosomes. To distinguish between these possibilities, we simultaneously imaged SsdA puncta and peroxisomes, marked by PexK-GFP (Salogiannis et al., 2016). SsdA puncta and peroxisomes displayed clearly distinct transport behaviors: most peroxisomes were stationary and moved only transiently, whereas SsdA puncta were persistently motile (**Fig. 4C, D**). Nevertheless, when peroxisomes did move, approximately 70% of movements coincided with detectable SsdA signal (**Fig. S4E**).

Together, these observations indicate that SsdA puncta can move independently of peroxisomes, using machinery in common with peroxisomes—PxdA and DipA—to hitchhike on early endosomes **(Fig. 4E**). In addition, our evidence suggests that an SsdA punctum and a peroxisome can hitchhike on the same early endosome.

### SsdA puncta are depleted near hyphal tips

Despite the abundance of early endosomes near hyphal tips, SsdA puncta were largely depleted in the most tip-proximal regions (**Fig. 3D; Fig. S3E; Video S4**). To quantify depletion, we examined SsdA-mScarlet3 and 3xTagGFP2-RabA puncta in 10 µm-long segments within the 50-µm region closest to the hyphal tip, in 132 hyphae **(Fig. 5A)**. Within the segment 10–0 µm from the tip, we observed a ∼2.5-fold decrease in puncta abundance compared to segments 50–40 µm and 40–30 µm from the tip (**Fig. 5B**). By contrast, early endosome abundance showed an opposite pattern, with highest abundance in the most tip-proximal segments (**Fig. 5B**). Interestingly, fluorescence of individual SsdA puncta also decreased in the most tip-proximal segments, while the fluorescence of RabA puncta remained constant (**Fig. 5C**). The combined effect of decreased SsdA puncta number and decreased fluorescence of individual SsdA puncta led to a 4-fold decrease in total SsdA punctate signal in the segment 10–0 µm from the tip compared to segments 50–40 µm and 40–30 µm from the tip (**Fig. S5A**). These results indicate that *A. nidulans* hyphae normally have an SsdA puncta depletion zone beginning at the hyphal tip, but they do not have a comparable RabA puncta depletion zone.

**Figure 5.**
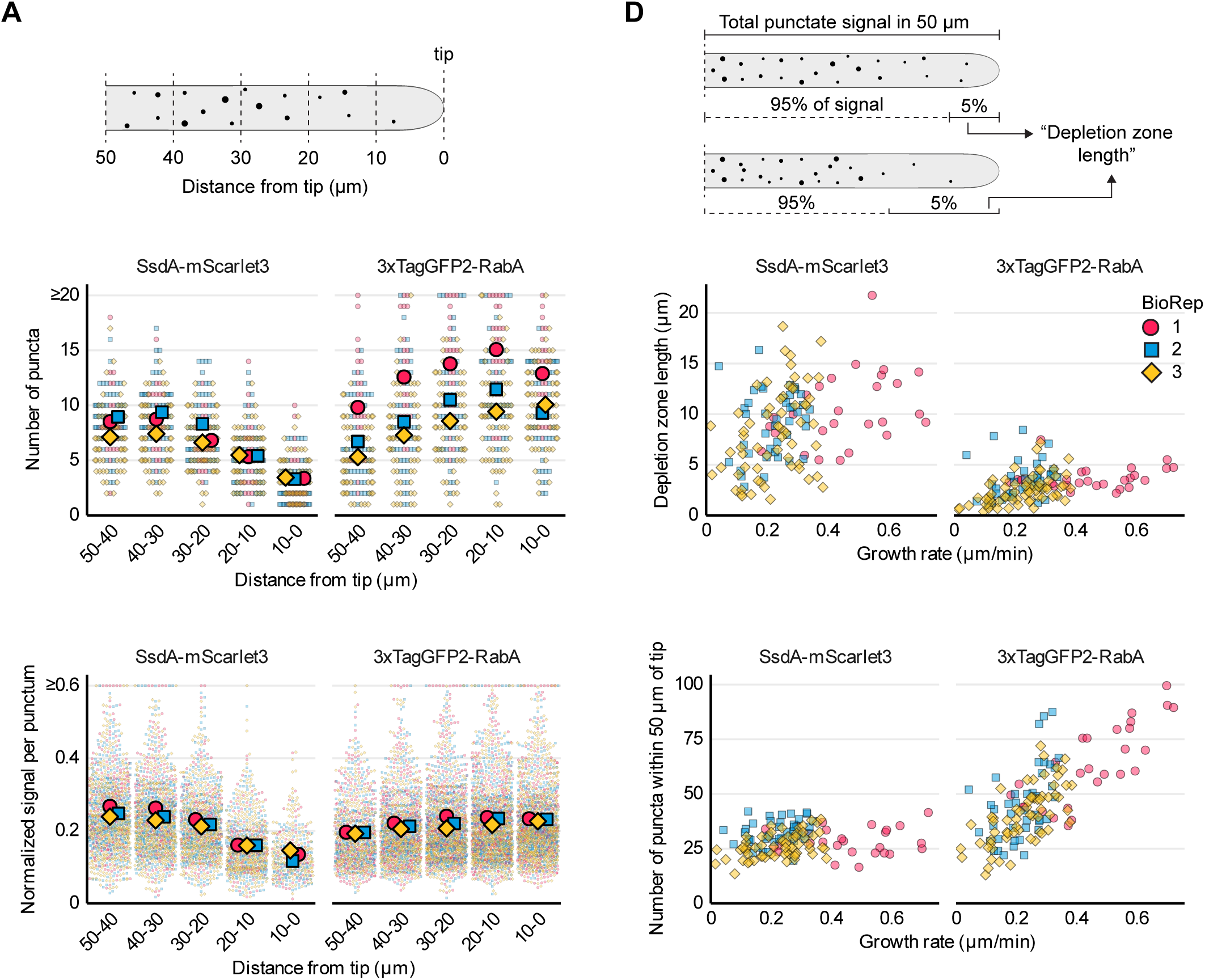
Decreased number and signal of SsdA puncta in regions closest to hyphal tips. **(A)** Schematic of distance-based quantification of puncta relative to the hyphal tip shown in (B) and (C); see main text for details. **(B)** Numbers of SsdA-mScarlet3 and 3xTagGFP2-RabA puncta in the 10-µm segments described in (A). Small symbols indicate individual hyphae; large symbols indicate means of biological replicates (BioReps). Different shapes/colors indicate distinct BioReps. **(C)** Fluorescent signal per punctum for SsdA-mScarlet3 and 3xTagGFP2-RabA in the 10-µm segments described in (A). Values were normalized to the minimum and maximum punctum signal within each BioRep. Small symbols indicate individual puncta; large symbols indicate means of BioReps. Different shapes/colors indicate distinct BioReps. **(D)** Schematic illustrating how length of SsdA puncta depletion zones was calculated; see main text for details. **(E)** SsdA puncta depletion zone length of SsdA-mScarlet3 or 3xTagGFP2-RabA versus hyphal growth rate. Symbols indicate individual hyphae. Different shapes/colors indicate distinct BioReps. **(F)** Total number of SsdA-mScarlet3 or 3xTagGFP2-RabA puncta in the 50-µm region closest to hyphal tips, plotted against hyphal growth rate. Symbols indicate individual hyphae. Different shapes/colors indicate distinct BioReps. Numbers of hyphae analyzed were 27, 42, and 63 for BioRep 1, 2, and 3 respectively. Growth rates were calculated from two timepoints (T0 and T1; see Methods). Panels B and C use data from T0 only. Y-axes in panels E and F represent mean values from T0 and T1. Data from T0 were also used for the quantification of SsdA colocalization with RabA shown in Fig. S3F, G.

### SsdA puncta depletion correlates with hyphal growth rate

We hypothesized that the SsdA puncta depletion zone might be related to hyphal growth. We first asked whether the length of a depletion zone correlates with growth rate. We defined the length of a puncta depletion zone as the length of the region (beginning at the hyphal tip) that contains 5% of the total punctate signal present within 50 µm of the hyphal tip (**Fig. 5D**). We found that faster-growing hyphae exhibited longer SsdA depletion zones, showing a moderate correlation (Spearman’s ρ = 0.39, p < 0.001; **Fig. 5E**). By contrast, when the same analysis was applied to RabA puncta, we did not observe a comparable magnitude of puncta depletion with growth rate (**Fig. 5E**). We next asked whether total puncta numbers within 50 µm of the hyphal tip also scaled with growth rate. Interestingly, numbers of RabA puncta increased strongly with increased growth rate (Spearman’s ρ = 0.71, p < 0.001), while numbers of SsdA puncta remained constant (**Fig. 5F**). Total punctate signal showed the same pattern (**Fig. S5B**). We conclude that the area number density of early endosomes scales linearly with growth rate, but the area number density of SsdA puncta does not increase proportionally. This disproportionality could reflect either active depletion of SsdA from tip-proximal regions or passive dilution of a fixed SsdA pool relative to an expanding endosome population. Finally, we asked whether hitchhiking contributes to the establishment of the SsdA puncta depletion zone. Deletion of either *pxdA* or *dipA* did not abolish the tip-proximal depletion of SsdA puncta (**Fig. S5C**), indicating that the depletion zone is established independently of endosomal hitchhiking.

### CotA kinase activity regulates SsdA puncta near hyphal tips

In fungi, NDR kinases have multiple roles in regulating polarized growth (Yarden et al., 1992; Maerz and Seiler, 2010). The *A. nidulans* NDR kinase CotA is essential for normal polarized growth and is enriched at hyphal tips (Shi et al., 2008). In *S. cerevisiae*, the CotA ortholog Cbk1 phosphorylates multiple NDR kinase consensus sites within the disordered N-terminus of Ssd1, regulating its function (Jansen et al., 2009; Kurischko et al., 2011a), and similar consensus sites are also present in SsdA (**Fig. 1A**). We therefore investigated whether the *A. nidulans* Cbk1 ortholog CotA might regulate SsdA localization near hyphal tips. Overexpressed CotA has been shown to localize to hyphal tips (Shi et al., 2008), but CotA expressed at endogenous levels in *Aspergillus fumigatus* was found to be undetectable (Martin-Vicente et al., 2024). Because CotA interacts with its coactivator MobB (Shi et al., 2008; **Fig. 6A**), we used MobB as a marker for potential CotA activity.

**Figure 6.**
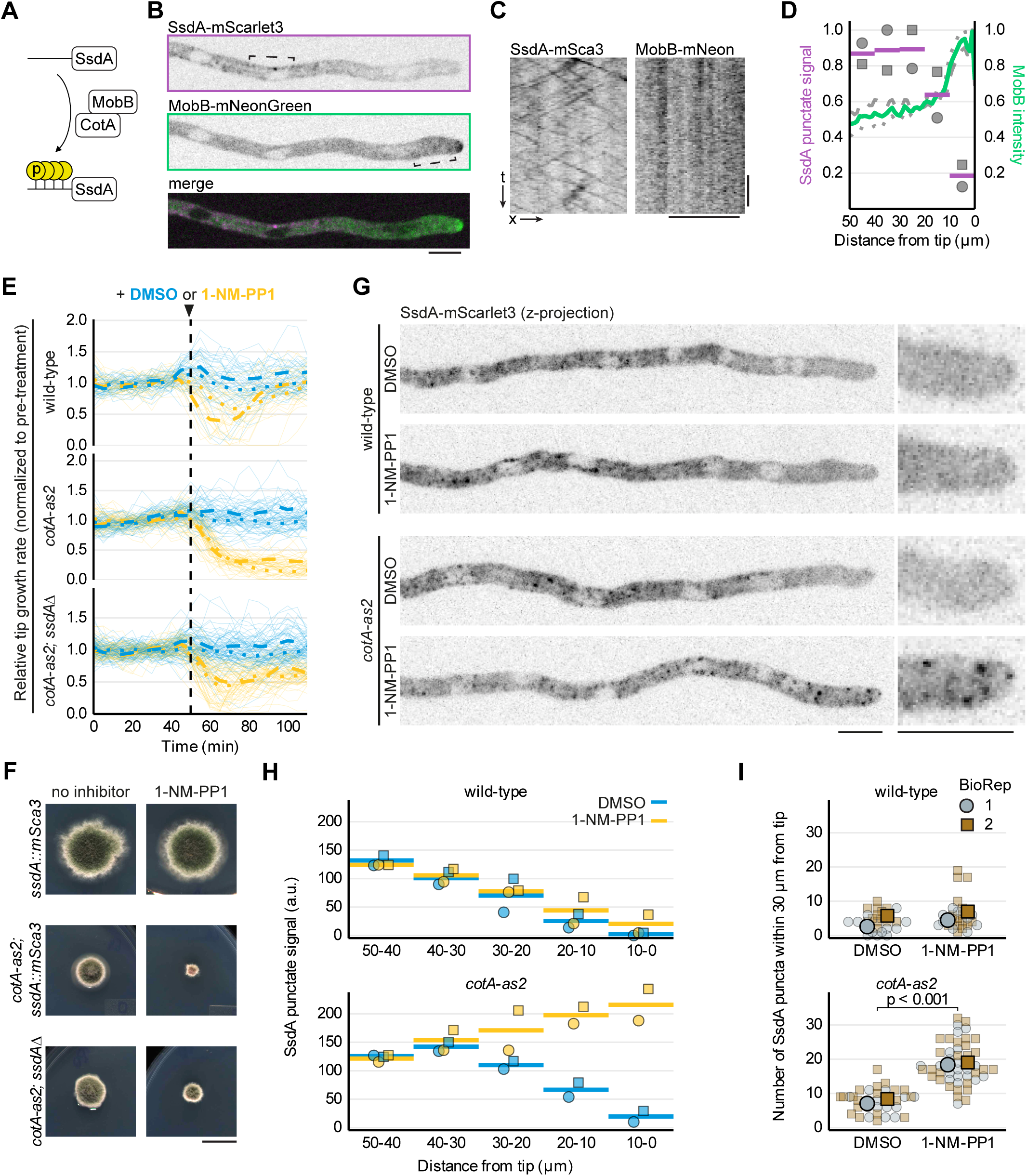
CotA kinase activity regulates hyphal growth rates and SsdA puncta localization. **(A)** Diagram showing predicted SsdA phosphorylation by NDR kinase CotA with its coactivator MobB, acting on 6 NDR kinase consensus sites in the disordered SsdA N-terminus (see Fig. 1A). **(B, C)** Zero timepoint from single-z-section near-simultaneous dual-channel movie of hypha expressing SsdA-mScarlet3 and MobB-mNeonGreen (B), with accompanying kymographs (C). Brackets in (B) indicate regions kymographed in (C). Faint MobB-mNeonGreen puncta were not detectably motile during the observation window, while SsdA-mScarlet3 puncta exhibited clear motility. **(D)** SsdA-mScarlet3 mean punctate signals and MobB-mNeonGreen intensity within the 50-µm region closest to hyphal tips, in hyphae expressing both proteins. SsdA punctate signal (sum of individual puncta signals) was quantified in discrete 10-µm segments, from maximum intensity z-projections. MobB intensity was measured continuously as a function of distance from the tip, from sum projections (see Methods). Gray symbols (SsdA) and broken lines (MobB) indicate individual biological replicates (BioReps). Magenta bars (SsdA) and green line (MobB) indicate means of BioReps. Values were normalized to the maximum value within each BioRep (maximum = 1). Number of hyphae: 24 per BioRep. **(E)** Changes in hyphal tip growth rates upon acute inhibition of kinase CotA. Growth rates are shown for wild-type, *cotA-as2* (analog-sensitive CotA), and *cotA-as2; ssdAΔ* strains before and after growth-medium exchange to fresh medium containing either vehicle (DMSO) or analog-sensitive kinase inhibitor 1-NM-PP1 (5 µM). Vertical line indicates time of medium exchange. Plotted traces are three-point moving averages of raw measurements, normalized to the mean growth rate before treatment for each hypha. Blue and yellow lines indicate DMSO- and 1-NM-PP1-treated hyphae, respectively. Thin lines indicate individual hyphae; thick dotted and dashed lines indicate means within each BioRep. Numbers of hyphae: wild-type/DMSO = 18, 29; wild-type/1-NM-PP1 = 12, 21; *cotA-as2*/DMSO = 27, 35; *cotA-as2*/1-NM-PP1 = 25, 29; *cotA-as2 ssdAΔ*/DMSO = 33, 45; *cotA-as2 ssdAΔ*/1-NM-PP1 = 25, 41 for BioReps 1, 2 respectively. **(F)** Colony phenotypes for the indicated genotypes, grown on minimal medium for 6 days at 25°C either in the absence of inhibitor or in the presence of 5 µM 1-NM-PP1. Panels shown here are also presented within a wider concentration/temperature range in Fig. S6C. Mutant *cotA-as2* grown on 5 µM 1-NM-PP1 exhibits significant growth inhibition and sporulation defects; these are partially rescued by deletion of *ssdA*. **(G)** Representative maximum-intensity z-projection micrographs of hyphae expressing SsdA-mScarlet3 in wildtype and *cotA-as2* backgrounds, 10 minutes after exchange to fresh medium containing either DMSO or 5 µM 1-NM-PP1. Expanded views of hyphal tips are shown at right. **(H)** Mean SsdA-mScarlet3 punctate signals in 10 µm-long segments within the 50-µm region closest to hyphal tips, for the four conditions shown in (G). Colors are as in (E). Symbols indicate means of hyphae within each BioRep; bars indicate means of BioReps. Values are arbitrary units (a.u.). Numbers of hyphae: wild-type/DMSO = 13, 11; wild-type/1-NM-PP1 = 10, 16; *cotA-as2*/DMSO = 15, 18; *cotA-as2*/1-NM-PP1 = 21, 25 for BioReps 1, 2 respectively. **(I)** Numbers of SsdA-mScarlet3 puncta within 30 µm of the hyphal tip. The same dataset was analyzed as in (G, H). Small symbols indicate individual hyphae; large symbols indicate means within BioReps. Different shapes/colors indicate distinct BioReps. Significance brackets indicate thresholds met in both BioReps (unpaired Welch’s t-test, two-tailed, per BioRep). Scale bars = 5 µm for micrographs, 1 cm for colony pictures.

We examined whether MobB localizes to the region in which SsdA puncta are depleted. MobB-mNeonGreen expressed from the endogenous locus was enriched at hyphal tips, precisely where SsdA puncta are depleted (**Fig. 6B, D**). Faint MobB-mNeonGreen puncta occasionally appeared elsewhere near tips but were not detectably motile, unlike SsdA puncta in the same observation window (**Fig. 6C**). We quantified MobB hyphal tip enrichment as the percentage of MobB signal in the tip-proximal 5 μm relative to the total MobB signal within 50 μm of the tip. We found no clear relationship between MobB hyphal tip enrichment and the length of the SsdA depletion zone (**Fig. S6A**), indicating that the depletion zone length is not simply determined by absolute MobB levels.

To test directly whether CotA kinase activity regulates SsdA distribution, we generated an analog-sensitive allele of CotA (M281A; *cotA-as2*), equivalent to alleles used for other Cbk1 orthologs (Weiss et al., 2002; Das et al., 2009; Kodama et al., 2017; Tay et al., 2018). This allele enables acute, specific inhibition of CotA using the ATP-competitive analog-sensitive kinase inhibitor 1-NM-PP1 (Bishop et al., 2000).

In the *cotA-as2* mutant, treatment with 1-NM-PP1 led to near-complete arrest of hyphal tip growth within approximately 10 minutes (**Fig. 6E; Fig. S6B**). By contrast, in wild-type hyphae, tip growth rates decreased transiently but recovered quickly (**Fig. 6E; Fig. S6B**). Both wild-type and *cotA-as2* hyphae grew normally when treated with DMSO (vehicle control; **Fig. 6E; Fig. S6B**).

To test whether the effects of CotA inhibition on hyphal tip growth are mediated through SsdA, we analyzed a *cotA-as2 ssdAΔ* double mutant (**Fig. 6E; Fig. S6B**). In the double mutant, 1-NM-PP1 treatment led to an approximately 35% decrease in tip growth rate, compared to the ∼80% decrease observed in the *cotA-as2* single mutant. Although CotA likely phosphorylates multiple substrates involved in growth, the partial suppression of decreased growth by *ssdAΔ* suggests that regulation of hyphal tip growth by CotA is mediated at least in part via *ssdA*.

Consistent with inhibition of CotA affecting hyphal tip growth, *cotA-as2* colonies exhibited dose-dependent growth defects on solid media in the presence of 1-NM-PP1 (**Fig. 6F; Fig. S6C**). Even without inhibitor, *cotA-as2* colonies were slightly more compact than wild-type, and this was exacerbated at higher temperatures (**Fig. S6C**). Inhibition of CotA also blocked sporulation and led to hyperbranching at early stages of colony development (**Fig. S6D**), reminiscent of the *cotA* deletion mutant (De Souza et al., 2013). Interestingly, as with tip growth, many of the colony phenotypes associated with CotA inhibition were partially suppressed by *ssdAΔ* (**Fig. 6F, S6C**).

We next asked whether acute inhibition of CotA affects SsdA localization. Upon DMSO treatment of wild-type and *cotA-as2* hyphae, or 1-NM-PP1 treatment of wild-type hyphae, SsdA puncta remained depleted in tip-proximal regions (**Fig. 6G**). However, within 10 minutes of 1-NM-PP1 treatment of *cotA-as2* hyphae, SsdA puncta appeared in the immediate vicinity of hyphal tips (**Fig. 6G**). Further analysis showed that 1-NM-PP1 had no effect on overall SsdA distribution in wild-type hyphae (**Fig. 6G–I**). By contrast, upon 1-NM-PP1 treatment of *cotA-as2* hyphae, the SsdA puncta depletion zone was completely abolished (**Fig. 6H**), and the total number of SsdA puncta within 30 μm of the hyphal tip increased approximately 2-fold (**Fig. 6I**). We conclude that CotA kinase activity is required to maintain the SsdA puncta depletion zone.

### Mutation of predicted CotA phosphorylation sites alters SsdA localization and colony morphology

To test whether CotA-mediated phosphorylation regulates SsdA localization and function, we generated mScarlet3-tagged SsdA mutants, expressed at the *ssdA* locus. We mutated all six predicted CotA phosphorylation sites in the disordered N-terminus to either alanine (SsdA-6A; nonphosphorylatable) or aspartic acid (SsdA-6D; phosphomimetic). We also generated a mutant lacking the 429 amino-acid disordered N-terminus (SsdA-NtermΔ). Western blotting confirmed that SsdA-NtermΔ and SsdA-6D were expressed at levels comparable to wild-type SsdA (**Fig. S7A, B**). We were unable to assess SsdA-6A protein levels because severe sporulation defects in this strain (see below) prevented the generation of sufficient biomass for protein extraction.

Live-cell imaging revealed striking differences in SsdA localization among the mutant strains (**Fig. 7A**). Both SsdA-6D and SsdA-NtermΔ displayed diffuse cytoplasmic distributions, with a marked decrease in the area number density of puncta compared to wild-type SsdA (**Fig. 7A, C**). By contrast, SsdA-6A puncta were readily visible throughout hyphae, with no apparent depletion zone (**Fig. 7A, D**), similar to wild-type SsdA upon CotA-as2 inhibition (**Fig. 6G, H)**. SsdA-6A puncta remained motile and were approximately 20% brighter than wild type (**Fig. S7C, D**). This indicates that N-terminal phosphorylation is not required for SsdA puncta motility but may influence the amount of SsdA associated with motile puncta.

**Figure 7.**
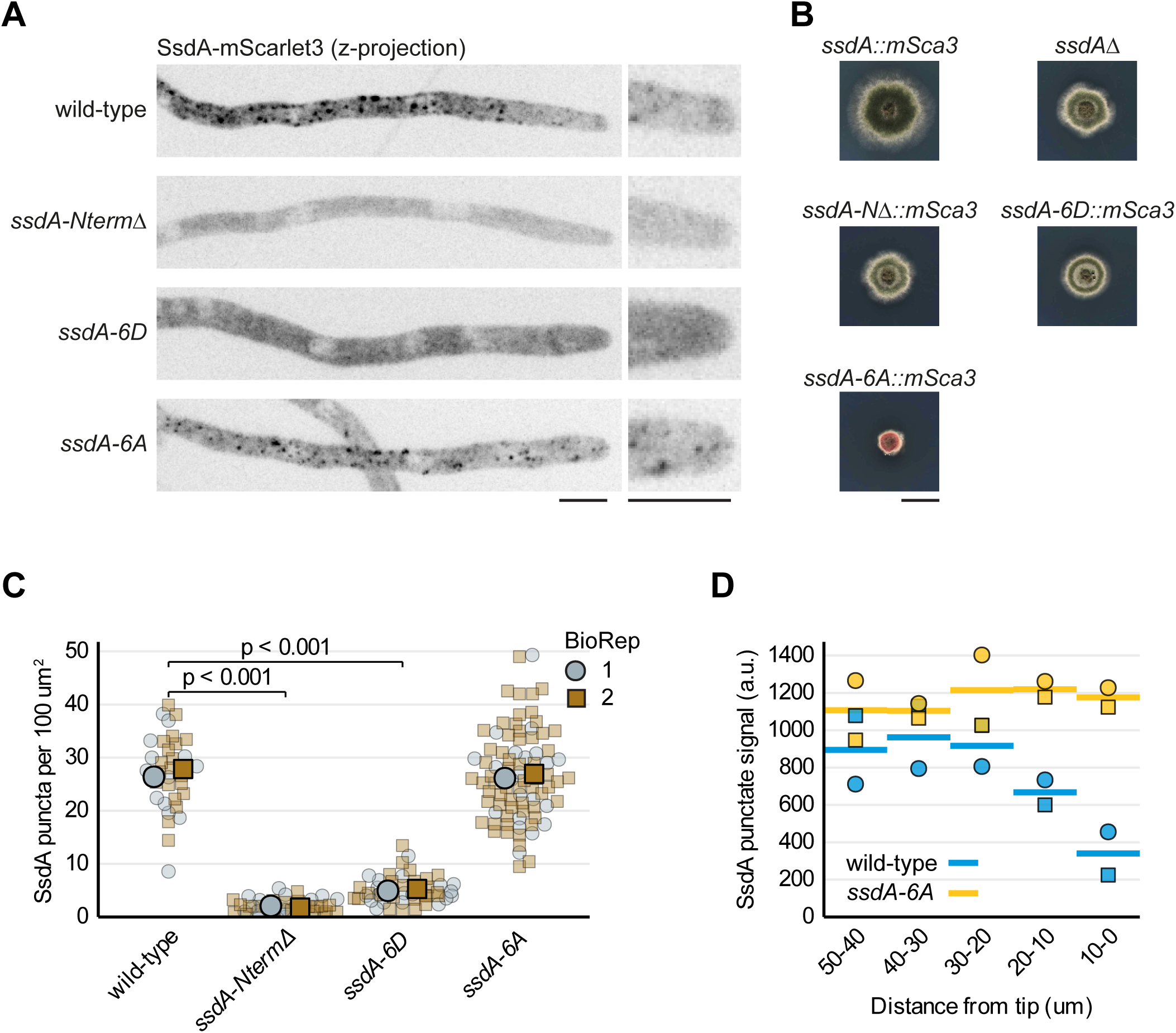
Mutation of predicted SsdA phosphorylation sites alters SsdA localization. **(A)** Maximum-intensity z-projection micrographs of hyphae expressing wild-type SsdA-mScarlet3 or mScarlet3-tagged SsdA N-terminal mutants (aa2-429Δ truncation mutant SsdA-NtermΔ, nonphosphorylatable mutant SsdA-6A, and phosphomimetic mutant SsdA-6D). Expanded views of hyphal tips are shown at right. **(B)** Colony phenotypes for wild-type and N-terminal mutants of mScarlet3-tagged *ssdA*, together with *ssdAΔ*, grown on minimal medium for 3 days at 30°C. The *ssdA-CA* mutant phenotype resembles *cotA-as2* grown on 1-NM-PP1 (see Fig. 6F). **(C)** SsdA-mScarlet3 puncta area number density in wild-type hyphae and the indicated *ssdA* N-terminal mutants. Small symbols indicate individual hyphae; large symbols indicate means within biological replicates (BioReps). Different shapes/colors indicate distinct BioReps. Significance brackets indicate thresholds met in both BioReps (Mann–Whitney *U*, two-tailed, per BioRep). Number of hyphae: wild-type = 14, 20; *ssdA-NtermΔ* = 21, 22; *ssdA-CD* = 20, 27; *ssdA-CA* = 22, 61 for BioReps 1 and 2, respectively. **(D)** SsdA-mScarlet3 mean punctate signals (from sums of individual puncta signals) in 10 µm-long segments within the 50-µm region of hyphae closest to hyphal tips, for wild-type and *ssdA-CA* strains. Small symbols indicate means of hyphae within each BioRep; bars indicate means of BioReps. Different shapes/colors indicate distinct BioReps. Values are arbitrary units (a.u.). Quantified from the same dataset as (C). Number of hyphae: wild-type = 14, 20; *ssdA-CA* = 7, 28 for BioReps 1 and 2, respectively. Scale bars = 5 µm for micrographs, 1 cm for colony pictures.

The *ssdA::mScarlet3* mutant strains also exhibited distinct colony morphologies (**Fig. 7B**). Colonies of *ssdA-6D* and *ssdA-NtermΔ* were slightly smaller than wild type, with more aerial hyphae and a sharp colony edge, recapitulating *ssdAΔ* colony morphology (**Fig. 7B**). By contrast, *ssdA-6A* colonies displayed a more severe phenotype, with morphology and sporulation defects closely resembling that of 1-NM-PP1-inhibited *cotA-as2* (**Fig. 7B**; compare to **Fig. 6F**). Early-stage *ssdA-6A* colonies were compact and hyperbranched, again similar to inhibited *cotA-as2* (**Fig. S7E**; compare to **Fig. S6D**).

Together, these data indicate that CotA-mediated phosphorylation of the SsdA N-terminus regulates SsdA puncta formation and is required for normal hyphal growth.

## Discussion

Here we have shown that SsdA, the *A. nidulans* ortholog of the *S. cerevisiae* RBP and proposed translational repressor Ssd1, is present in normal growing hyphae as cytoplasmic puncta that represent mRNPs (SsdA-mRNPs). In wild-type hyphae, many SsdA puncta move along microtubules, in both anterograde and retrograde directions, via hitchhiking on early endosomes. SsdA hitchhiking requires endosomal adaptor proteins PxdA and DipA, which have previously been associated only with peroxisome hitchhiking on endosomes. Unexpectedly, SsdA puncta are largely depleted in hyphal tip-proximal regions, and depletion depends on kinase activity of NDR kinase CotA: acute inhibition of analog-sensitive CotA causes rapid accumulation of SsdA puncta near tips, and an SsdA mutant predicted to be nonphosphorylatable by CotA displays the same accumulation of puncta near hyphal tips, together with severe growth defects. Collectively, our findings establish SsdA-mRNPs as a new class of cargo for the endosomal hitchhiking system in *A. nidulans* and reveal that NDR kinase signaling can spatially regulate SsdA-mRNP distribution during polarized growth. Below, we discuss the nature of SsdA puncta, their mode of hitchhiking and their regulation by CotA, as well as the potential biological functions of SsdA-mRNP dynamics in the context of a hypothesized role for SsdA in translational regulation.

### SsdA puncta and mRNP complexes

SsdA puncta likely encompass several different states of SsdA: 1) SsdA-mRNPs hitchhiking on early endosomes; 2) SsdA-mRNPs not associated with endosomes; and 3) SsdA not associated with mRNA. Our results further suggest that multiple factors contribute to puncta formation.

One key factor is mRNA binding. Our photobleaching experiments showing colocalization of poly(A)-binding protein FabM with motile SsdA puncta in wild-type hyphae support the conclusion that motile SsdA puncta represent SsdA-mRNP complexes. In the *ssdA-WAKA* mutant, which is predicted to exhibit strongly decreased RNA binding (Bayne et al., 2022), numbers of SsdA puncta in hyphae are ∼3-fold lower than in wild-type hyphae. In addition, puncta in *ssdA-WAKA* have a lower fluorescent signal (see Discussion about puncta signal below). This analysis of puncta in *ssdA-WAKA* provides the strongest experimental evidence that most SsdA-puncta in wild-type hyphae represent SsdA-mRNP complexes, whether or not they are motile or associated with endosomes. We note that SsdA-mRNP puncta could contain a single mRNA bound to multiple SsdA molecules or multiple mRNAs bound to one or more SsdA molecules.

The SsdA N-terminus is also critical for puncta formation. We detected almost no puncta in the *ssdA-NtermΔ* mutant, even though SsdA-NtermΔ protein is expressed at wild-type levels. Moreover, the few SsdA-NtermΔ puncta that are detectable are extremely faint. Currently it is unclear how the SsdA N-terminus contributes to SsdA puncta formation. One possibility is that the N-terminus, which is predicted to be unstructured, leads to aggregation of SsdA, either through homomeric or heteromeric interactions, independent of mRNA binding. Under stress, *S. cerevisiae* Ssd1 can appear in foci that localize with P-bodies (Jansen et al., 2009; Kurischko et al., 2011b; Kurischko and Broach, 2017; Gutierrez et al., 2026). In addition, overexpressed Ssd1 N-terminal fragments have been observed in foci (Kurischko et al., 2011a, 2011b).

Although budding yeast Ssd1 lacking the N-terminus can bind cognate RNA *in vitro* with low-nanomolar affinity (Bayne et al., 2022), the SsdA N-terminus could also potentially contribute to mRNA binding *in vivo*. For example, aggregation of SsdA via its N-terminus might enable even stronger, cooperative, binding *in vivo* to target mRNAs containing multiple SsdA binding sites.

Association with early endosomes also seems to enhance SsdA puncta formation and/or fluorescent signal. SsdA puncta in *pxdAΔ* and *dipAΔ* mutant hyphae have lower signal and are fewer in number than in wild-type hyphae (albeit greater in number than in *ssdA-WAKA*), in addition to being non-motile. Consistent with this, in wild-type hyphae, SsdA puncta that colocalize with RabA endosomes tend to have a higher SsdA signal than puncta that do not colocalize with RabA. This correlation supports the idea that association with early endosomes may bring together multiple SsdA-mRNPs.

It is not clear how the SsdA puncta we observe here relate to puncta of Ssd1 homologs observed in other systems. In *S. cerevisiae*, Ssd1 foci are observed under conditions of stress or inhibition of Cbk1 (Jansen et al., 2009; Kurischko et al., 2011b; Kurischko and Broach, 2017; Gutierrez et al., 2026), while Ssd1 is normally diffuse in the cytosol in unstressed cells (Dubreuil et al., 2019). Similar observations have been made for the *Schizosaccharomyces pombe* SsdA ortholog Sts5 (Nuñez et al., 2016). By contrast, our data show that *A. nidulans* SsdA puncta are constitutive, motile, and abundant under normal vegetative conditions. Similar motile puncta have also been observed for the SsdA ortholog GUL1 in the filamentous ascomycete *S. macrospora* (Stein et al., 2020), raising the possibility that the mechanism of SsdA/GUL1 puncta assembly is conserved among filamentous ascomycete fungi (see below). Further experiments will be needed to test whether SsdA can also localize to P-bodies and under what conditions.

### SsdA endosomal hitchhiking

One of our key findings is that SsdA endosomal hitchhiking relies on the same adaptor proteins (PxdA and DipA) that are required for endosomal hitchhiking by peroxisomes and Woronin bodies (Salogiannis et al., 2016; Songster et al., 2023). It remains to be determined whether PxdA and DipA interact directly with SsdA-mRNP components or require additional bridging factors. The recent identification of *A. nidulans* AcbdA as a peroxisomal membrane-localized cargo receptor (Driscoll et al., 2025) suggests that PxdA and DipA may constitute a shared endosomal docking platform to which distinct cargoes bind via different receptors.

It is also unclear whether SsdA mediates association of its target mRNAs with endosomal hitchhiking machinery, or whether SsdA is primarily a passenger on mRNP complexes that can interact with hitchhiking machinery independently of SsdA. The presence of motile FabM (poly(A)-binding protein) puncta in *ssdAΔ* hyphae is consistent with either possibility; this reveals only that at least some mRNAs do not require SsdA for movement. Further elucidation of what role, if any, SsdA plays in recruitment of mRNAs to the hitchhiking machinery will depend on identification and imaging of specific hitchhiking mRNAs.

The role of PxdA and DipA in endosomal hitchhiking of mRNPs is unlikely to be restricted to *A. nidulans*. Prior to our work, the *S. macrospora* SsdA ortholog GUL1 was shown to hitchhike on early endosomes, in association with poly(A)-binding protein Pab1 (Stein et al., 2020). Similarly, aggregates of the *Neurospora crassa* SsdA ortholog, GUL-1, were observed to move in hyphae in a microtubule-dependent manner (Herold et al., 2019); this may also represent endosomal hitchhiking. Because both PxdA and DipA are conserved among filamentous ascomycetes (Pezizomycotina), including *S. macrospora* and *N. crassa* (Songster et al., 2023), we hypothesize that these two proteins may function more generally as multi-purpose adaptors for endosomal hitchhiking of multiple cellular components in Pezizomycotina species.

This accords with current ideas that early endosomes function as general-use transport platforms in filamentous fungi more broadly, although the specific molecular machinery used in different organisms can vary (Christensen and Reck-Peterson, 2022; Vázquez-Carrada et al., 2026). In the basidiomycete fungus *U. maydis*, where endosomal hitchhiking was first discovered (König et al., 2009; Baumann et al., 2012) early endosomes carry mRNPs, peroxisomes, lipid droplets, and ER (Baumann et al., 2012; Guimaraes et al., 2015). However, several of the key proteins involved in mRNP hitchhiking in *U. maydis*, such as RBP Rrm4 (Pohlmann et al., 2015) and poly(A)-binding protein interactor Upa2 (Jankowski et al., 2019) are not present in ascomycete fungi, despite their broad conservation in basidiomycetes (Müller et al., 2019). Conversely, PxdA is found only in Pezizomycotina species (Songster et al., 2023).

Association of mRNAs with motile membrane-bound organelles also occurs beyond fungi: in mammalian neurons, mRNAs can be transported on lysosomes, tethered by Annexin A11 (Liao et al., 2019), and on early endosomes, tethered by the FERRY complex (Schuhmacher et al., 2023).

### Spatial regulation of SsdA-mRNP puncta by NDR kinase CotA

Another of our key findings is that CotA kinase activity regulates the spatial distribution of SsdA puncta in hyphae. During normal hyphal growth, we observe an SsdA puncta depletion zone in tip-proximal regions. However, acute inhibition of CotA causes rapid accumulation of SsdA puncta in tip-proximal regions and greatly impaired hyphal tip growth. Furthermore, mutation of predicted consensus CotA phosphorylation sites within the SsdA N-terminus (*ssdA-6A*) recapitulates the effects of CotA inhibition on puncta, strongly suggesting that CotA acts on puncta directly via phosphorylation of the SsdA N-terminus.

How might N-terminal phosphorylation affect the properties of SsdA? Here we consider two plausible, non-exclusive scenarios. First, in line with a proposed role for the SsdA N-terminus in puncta formation (see above), phosphorylation may dissolve SsdA N-terminus-driven aggregates. Second, in accordance with the importance of mRNA binding for SsdA puncta formation, phosphorylation may negatively affect SsdA binding to target mRNAs. Both of these scenarios are consistent with the phenotype of the *ssdA-6D* mutant, in which puncta formation is severely compromised. We note that both scenarios are also agnostic as to whether SsdA target mRNAs would be released from endosomes as a result of SsdA N-terminus phosphorylation (see below and **Model Fig. 8**).

**Figure 8.**
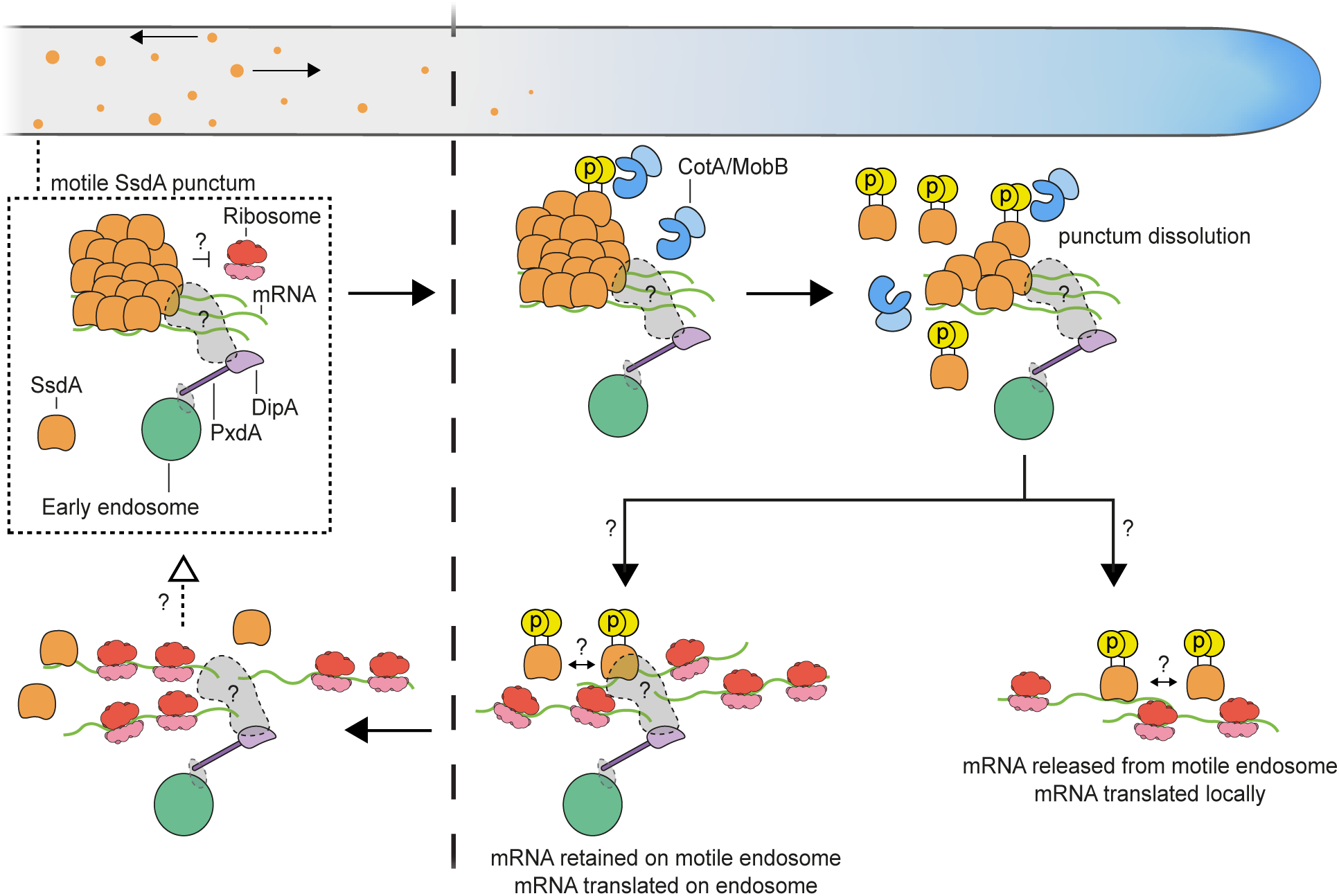
Speculative model for SsdA-mediated spatially restricted translational regulation near hyphal tips. Motile SsdA puncta represent SsdA-mRNP aggregates that hitchhike on early endosomes via adaptor proteins PxdA and DipA. Additional bridging factors (gray shapes) may be required for attachment of SsdA-mRNPs and/or PxdA/DipA to early endosomes. mRNAs bound by SsdA are likely to encode cell-wall and other polarity-related proteins (Fig. S1C) and are proposed to be translationally repressed within SsdA puncta. As endosomes carrying SsdA puncta approach the hyphal tip, the NDR kinase CotA (with its co-activator MobB) phosphorylates SsdA, leading to dissolution of puncta. Whether individual phosphorylated SsdA molecules remain associated with target mRNAs after dissolution is unknown. Following punctum dissolution, target mRNAs become competent for translation, as indicated by ribosome association. Two non-exclusive scenarios are depicted for the subsequent fate of derepressed mRNAs. In the first scenario (bottom right), mRNAs are released from early endosomes into the tip-proximal cytoplasm and are translated locally. In the second scenario (bottom centre), mRNAs remain associated with early endosomes after punctum dissolution and are translated on the endosomal surface; continued bidirectional endosome motility would distribute nascent polypeptides across the tip-proximal region (see main text). Question marks denote steps or interactions that remain to be experimentally tested.

Consistent with phosphorylation limiting punctum assembly, SsdA-6A puncta are modestly brighter than wild-type, indicating that more SsdA accumulates in puncta when phosphorylation is blocked. In *S. pombe*, phosphorylation of SsdA ortholog Sts5 by Orb6, the CotA ortholog, promotes binding by the 14-3-3 protein Rad24 and inhibits recruitment into P-bodies (Nuñez et al., 2016). Whether SsdA similarly interacts with 14-3-3 or other proteins upon phosphorylation is unknown, but this represents a plausible mechanism.

### Biological functions of SsdA puncta regulation

How might spatial regulation of SsdA puncta contribute to normal hyphal growth *in A. nidulans*? In *S. cerevisiae,* the SsdA ortholog Ssd1 is thought to function as a translational repressor that is negatively regulated via N-terminal phosphorylation by the CotA ortholog Cbk1 (Jansen et al., 2009; Hall and Wallace, 2022). While the mechanism by which Ssd1 represses translation of target mRNAs remains unclear, many target mRNAs encode proteins involved in cell polarity and cell wall biogenesis (Bayne et al., 2022). Furthermore, the nucleotide-sequence motifs used by Ssd1 to recognize target mRNAs in *S. cerevisiae* are also enriched in mRNAs encoding similar cell-polarity and cell-wall proteins in *A. nidulans*, *Candida albicans*, *S. pombe*, *A. fumigatus*, and *N. crassa* (Herold et al., 2019; Bayne et al., 2022; Schrettenbrunner et al., 2026). Together with the conservation of Ssd1 orthologs among all branches of Ascomycete fungi (Ballou et al., 2021), this strongly supports the idea that SsdA and other Ssd1 orthologs are RBPs with conserved target specificity and broadly conserved roles in regulating mRNAs related to cell polarity and cell-wall growth. From this perspective, it seems likely that SsdA-mRNP puncta hitchhiking on early endosomes contain target mRNAs (involved in cell wall/cell polarity) in a translationally repressed form, and that local derepression near hyphal tips, via CotA, is a critical feature of normal hyphal growth (**Fig. 8**).

In addition to being consistent with all our results involving perturbations to the SsdA-CotA system, this interpretation of SsdA puncta as translationally repressed mRNPs is also supported by correlative data in normal, non-perturbed hyphae. Specifically, in faster-growing hyphae, tip-proximal SsdA puncta depletion zones are longer than in slower-growing hyphae. Furthermore, while the total number of early endosomes within tip-proximal regions of hyphae increases linearly with hyphal growth rate—presumably to facilitate rapid growth—the total number of SsdA puncta does not increase with growth rate. Overall, this suggests a model in which repressive SsdA near hyphal tips may limit hyphal growth, and that faster hyphal growth may be associated with more efficient dissolution of SsdA puncta in hyphal tip-proximal regions, perhaps driven by increased CotA kinase activity. Interestingly, a relationship between hyphal growth rate and absence of repressive mRNP structures at hyphal tips has recently been described in the syncytial fungus *Ashbya gossypii* (Geisterfer et al., 2026).

A key question emerging from our work concerns the fate of SsdA target mRNAs as endosomes enter the SsdA puncta depletion zone. Within this region, SsdA puncta appear to dissolve, but the endosomes upon which they were hitchhiking remain present and continue to move bidirectionally. What happens to SsdA target mRNAs after SsdA puncta dissolution, and where are they translated?

We imagine two scenarios, with distinct consequences for spatial control of translation (**Fig. 8**). In the first scenario, dissolution of SsdA puncta is accompanied by release of SsdA target mRNAs from early endosomes into the tip-proximal cytoplasm. These mRNAs would then be translated close to the hyphal tip, positioning newly synthesized proteins where they might be needed most. In the second scenario, derepressed SsdA target mRNAs would be retained on early endosomes even after SsdA puncta dissolution. These mRNAs could then be translated on the surface of bidirectionally motile early endosomes, broadening the positioning of newly synthesized proteins (**Fig. 8**).

We favor the second scenario, primarily for theoretical reasons. Predicted SsdA targets are highly enriched for transcripts encoding plasma membrane and cell wall proteins, which generally require transit through the endoplasmic reticulum (ER) and Golgi apparatus before reaching the plasma membrane. In long hyphae of *A. nidulans*, the peripheral ER is enriched in a broad region extending ∼20–40 µm from the hyphal tip (Markina-Iñarrairaegui et al., 2013). An equivalent broadening of the distribution of derepressed mRNAs via endosomal hitchhiking could facilitate access of nascent polypeptides to the secretory machinery, particularly if the amount of available ER near the hyphal tip was limiting for secretion. This mode of translation could serve alongside a mechanism of indirect endocytic recycling, in which polarity proteins are maintained at hyphal tips by endocytosis, trafficking to the trans-Golgi network, and retargeting to hyphal tips (Hernández-González et al., 2018).

Our findings concerning the dynamics and regulation of SsdA puncta in *A. nidulans* add to a growing body of evidence that spatial regulation of RBPs is a fundamental feature of tip-growing fungal cells (Hall and Wallace, 2022; Vázquez-Carrada et al., 2026; Geisterfer et al., 2026).

## Methods

### Strains and growth conditions

*A. nidulans* strains used in this study are listed in **Table S1** (also available on Zenodo as .xlsx; doi: 10.5281/zenodo.19401665). Strains were grown on yeast glucose (YG) medium (20 g/L glucose, 5 g/L yeast extract, Hutner’s trace elements) or *A. nidulans* minimal medium (ANMM) as in Hill and Kafer (2001). Media were supplemented as required for auxotrophic markers with uridine (2.44 g/L), uracil (1.01 g/L), pyridoxine (0.5 g/L), and para-aminobenzoic acid (1 g/L).

### Molecular cloning

Plasmids and oligonucleotides used in this study are described in **Table S2** and **Table S3** (also available on Zenodo as .xlsx; doi: 10.5281/zenodo.19401665). All plasmids were assembled via Golden Gate (Engler et al., 2008) or Gibson (Gibson et al., 2009) cloning. Plasmid maps are available from Zenodo (doi: 10.5281/zenodo.17210183).

Where plasmids included synthetic gene fragments, these were codon optimized for *A. nidulans* as in Modaffari et al. (2024). Plasmid sequence was verified via Sanger sequencing (Genewiz, UK) or whole-plasmid sequencing (Plasmidsaurus, UK).

### Strain construction and verification

#### CRISPR-CasS genome editing

CRISPR-Cas9 genome editing enabled markerless genetic modifications using short homology arms (30-100 bp), preserving endogenous regulatory elements. We used the Golden Gate-based plasmid system from Modaffari et al. (2024).

sgRNAs targeting genomic loci were cloned into CRISPR-Cas9 vectors pDM026 (pyrG selection; Addgene #216808) or pDM068 (NAT selection; Addgene #216811) via Golden Gate cloning with PaqCI (New England Biolabs, Cat. No. R0745), following Modaffari et al. (2024). We verified correct spacer insertion by loss of GFP fluorescence in transformed *E. coli* and Sanger sequencing of recovered plasmids.

We generated homology-repair templates by PCR (for gene tagging) or oligonucleotide annealing (for point mutations). For fluorescent-protein tagging, templates contained 30-80 bp homology arms flanking codon-optimized mScarlet3 or mNeonGreen sequences (Addgene #216812-216815). The analog-sensitive *cotA-as2(M281A)* allele was generated by ends-in integration (see below), allowing subsequent marker excision.

Protoplasts were prepared as described by Oakley et al. (2012), with minor modifications, and co-transformed with 1–2 μg CRISPR plasmid and 0.5–1.5 μg homology-repair template, as described in Modaffari et al. (2024). Transformations were plated on YG agar with 1 M sucrose, supplemented with 200 μg/mL nourseothricin for NAT selection where needed. Transformation plates were incubated at 30°C for 1 day and then at 37°C for 3-4 days. Genotypes were verified by PCR and/or Sanger sequencing (Genewiz, UK), and/or microscopy visualization of fluorescent protein localization.

#### Strain construction with integrating selectable markers

Traditional homologous recombination with long (>1 kb) homology arms was used for gene deletions and constructions requiring stable marker integration. Gene deletions replaced entire coding sequences with *pyrG* (from plasmid pDM106) or *NAT* (from plasmid pDM145) markers.

Protoplast transformations used 2-5 μg linearized DNA, with selection maintained on YG agar with 1 M sucrose and appropriate supplements. Correct integration was verified by PCR and/or sequencing.

To construct *ssdA-NtermΔ* (aa2-429Δ), *ssdA-6A* (S26A, S126A, S183A, S255A, S300A, S339A), and *ssdA-6D* (S26D, S126D, S183D, S255D, S300D, S339D) strains, we generated plasmids (pDM159, pDM160, pDM161) containing full-length *ssdA::mScarlet3* coding sequences with the relevant mutations, plus flanking homology regions. Coding sequence plus homology regions were PCR-amplified and transformed as linear DNA into an *ssdAΔ::pyrG* deletion strain (aDM039). Recombinants were selected on YG medium containing 1 M sucrose + Uridine + Uracil + 0.1% 5-fluoroorotic acid (FOA) and verified by sequencing.

Ends-in homologous recombination, in which a linearized plasmid integrates via single crossover to create gene duplications flanking the marker, was used to generate the *cotA-as2::pyrG* allele (strain aDM062). Subsequent self-crossing and 5-FOA selection (YG + Uridine + Uracil + 0.15% 5-FOA) for loss (“loop-out”) of *pyrG* yielded a markerless *cotA-as2* strain (aDM064).

#### Genetic crosses

For genetic crosses, we followed the procedures described by Todd et al. (2007). Briefly, heterokaryons were established by co-inoculating parental strains on non-selective medium, followed by transfer of agar plugs containing hyphae from the zone where the mycelial fronts met onto appropriate selective medium. After cleistothecia formed, ascospores were plated on non-selective medium. Desired recombinants were verified by transfer to selective medium and/or by microscopy.

#### Strain purification

To obtain clonal isolates, strains were purified either by single-spore dissection or by successive rounds of streak purification. For single-spore dissection, sporulating colonies were briefly resuspended by pipetting 0.01% (v/v) Tween-80 over the colony surface, then the resulting spore suspension was spotted onto the lower region of a non-selective agar plate. Individual spores were separated by dissection in a regular grid (∼1.25 cm spacing) using an MSM 400 semi-automated tetrad dissection microscope (Singer Instruments, UK). To prevent cross-contamination, the initially inoculated spore spot was excised from the agar before incubation. For streak purification, spores were streaked to single colonies and re-streaked as needed until colonies of uniform morphology were obtained.

#### Genotyping PCR and DNA extraction

Genotypes were verified by PCR amplification of the relevant genomic locus directly from spores or from extracted genomic DNA. For spore PCR, lysates were prepared by heat/freeze treatment (1–10 µL of a medium-dark green spore suspension heated to 95°C for 15 min and then snap-frozen in liquid nitrogen or frozen at −70°C for 10 min) (Fraczek et al., 2019) or by mechanical disruption of spores by squashing a small volume of spore suspension (in 0.01% Tween-80) between a glass slide and a 22 x 22 mm No. 1.5 coverslip (Yuan et al., 2023). Routine genotyping used LongAmp Taq (New England Biolabs; M0323) or DreamTaq (Thermo Fisher Scientific; K1081); amplicons requiring sequence confirmation were generated with the high-fidelity Platinum SuperFi II polymerase (Thermo Fisher Scientific; 12361010). PCR products were purified and verified by Sanger sequencing (Genewiz, UK) or whole-amplicon sequencing (Plasmidsaurus, USA).

When higher-quality PCR template was required, we extracted genomic DNA from spores using a CTAB-based protocol adapted from Fraczek et al. (2013). Briefly, spores were harvested and pelleted, resuspended in CTAB extraction buffer (2% CTAB, 100 mM Tris-HCl pH 8.0, 1.4 M NaCl, 10 mM EDTA), mechanically disrupted with 0.5 mm glass beads, and incubated at 65°C. After RNase A treatment, DNA was extracted with chloroform:isoamyl alcohol (24:1), precipitated with isopropanol, washed with 70% ethanol, and resuspended in water for downstream applications.

### Protein extraction and western blot

Denaturing protein extraction was performed by adapting a trichloroacetic acid (TCA) protocol (Grallert and Hagan, 2017). Approximately 1 x 10^8^ conidia were grown in 50 mL ANMM with appropriate supplements for 16 hours at 30°C. The culture was then harvested by filtering through Miracloth (Millipore #475855), and excess liquid was removed by placing the Miracloth (containing wet mass) within a Germinated Aspergillus Recovery Lever Inducing Compaction (GARLIC) Press (Amazon UK) and applying a moderate-to-firm squeeze. Biomass was then flash-frozen in liquid nitrogen. Mycelia (approximately 100 mg wet weight) were treated with 500 µL ice-cold 20% TCA and 1 mL of 0.5 mm zirconium beads in 2 mL screwcap tubes. Cells were disrupted using a FastPrep-24 (MP Biomedicals, USA) bead beater at power 4.5 for two 15-second cycles, with 2 minutes of cooling on wet ice between cycles. After adding 400 µL of ice-cold 5% TCA, the bottom of the tubes was pierced with a hot needle. The pierced tubes were then placed within 5 mL polypropylene tubes and centrifuged at 4,000 rpm for 3 minutes at 4°C to separate the lysate from the beads. The lysate was then transferred to fresh 1.5 mL tubes and centrifuged at 13,000 rpm for 6 minutes at 4°C. Following removal of the supernatant, the resulting pellet was washed once with 1.5 M Tris pH 8.8 to neutralize the acidic pH and improve protein solubilization (Todd et al., 2005). Pellets were resuspended in Laemmli sample buffer containing 5% mercaptoethanol, boiled for 5 minutes at 95°C, and centrifuged at 13,000 rpm for 15 minutes at room temperature. The supernatant was transferred to a new tube. Final protein concentration was measured by Cydex Blue protein assay (Rabilloud, 2018), which is compatible with both SDS and reducing agents.

For SDS-PAGE and western blotting, approximately 40 µg of protein per lane was resolved on a 3–8% Tris-Acetate gel and transferred to a 0.2 µm nitrocellulose membrane. Total protein (used for normalization of western blot signals) was visualized with REVERT 700 Total Protein Stain (LI-COR). Membranes were probed for mScarlet3 using a primary goat anti-mScarlet antibody (St. Johns Laboratory, STJ140286; 1:6,000) and an IRDye 800CW Donkey anti-Goat secondary antibody (LI-COR, 926-32214; 1:20,000). Western blots were quantified using Image Studio software (LI-COR).

### Microscopy

Spinning-disk confocal imaging was performed on Nikon Ti2-E inverted microscopes equipped with Yokogawa CSU-W1 scanner units and a piezoelectric Z device for z-position control. Imaging was performed in temperature-controlled rooms (typically 18– 22°C). The first system was equipped with two Photometrics Prime 95B cameras and an Ohm Medical Systems FRAP/ablation unit and was used with either a Plan Apo VC (100x/1.4 NA) or Plan Apo Lambda D (100x/1.45 NA) oil-immersion objective and either a tri-band (Di01-T405/488/561; Semrock) or quad-band (Di01-T405/488/568/647; Semrock) dichroic. For this system, green fluorophores were excited with a 488 nm laser and detected with a 550 nm band-pass filter; red fluorophores were excited with a 561 nm laser and detected with a 568 nm long-pass filter. A second Ti2-E/CSU-W1 system was equipped with a single Photometrics Prime 95B camera with a quad-band dichroic (Di01-T405/488/568/647; Semrock) and was used with either a Plan Apo TIRF (100x/1.49 NA) oil-immersion objective or Plan Apo Lambda (40x/0.95 NA) dry objective. For this system, green fluorophores were detected with a 525/30 band-pass filter and red fluorophores with a 600/52 band-pass filter.

Images were acquired either as single z-sections or as z-stacks containing 11 sections with 0.6 µm spacing. For dual-channel single z-sections involving SsdA-mScarlet3 with PexK-GFP and SsdA-mScarlet3 with MobB-mNeonGreen, channels were acquired sequentially. For all other dual-channel single z-section imaging, channels were acquired simultaneously, using the two-camera system. For dual-channel z-stacks involving SsdA-mScarlet3 with MobB-mNeonGreen, channels were acquired sequentially, collecting a full z-stack in one channel followed by a full z-stack in the other channel; for dual-channel z-stacks involving SsdA-mScarlet3 and 3 xTagGFP2-RabA, channels were acquired simultaneously.

Early-stage colony micrographs were acquired on a widefield Nikon Ti2-E microscope.

### Live-cell imaging and drug treatments

For live-cell imaging experiments, spores freshly harvested from 2–3-day old colonies were inoculated onto ANMM plates containing 1% agarose and supplements as required, then incubated at 25–32 °C for 16–24 h. Agar plugs containing actively growing hyphae were excised from the plates and inverted onto glass-bottom Ibidi 4-well µ-Slide chambers (Cat. No. 80427; Ibidi GmbH, Germany). For experiments requiring medium exchange, 2 x 10⁴ spores were inoculated in 700 µL of liquid ANMM directly into the Ibidi 4-well chambers, followed by overnight incubation under the same conditions. Imaging was performed in the same chambers. Prior to image acquisition, samples were equilibrated to the imaging temperature (18–22 °C, room-controlled) by placement on the microscope stage for at least 30 min.

For medium exchange during imaging, the existing medium was gently removed using a 1 mL transfer pipette (Fisher Scientific, Cat. No. 13469118) and replaced with fresh medium containing the indicated drugs or vehicle control directly on the microscope stage. Replacement media were pre-equilibrated to the microscope room temperature prior to addition.

Methyl benzimidazol-2-yl carbamate (MBC) was dissolved in DMSO to prepare a 2.5 mg/mL stock solution. Working medium containing 25 µg/mL MBC was prepared fresh alongside a DMSO-only control (1% final DMSO concentration in both cases). Medium was exchanged as described above, and z-stacks were acquired 10 minutes after exchange.

The analog-sensitive kinase inhibitor 1-(tert-butyl)-3-(naphthalen-1-ylmethyl)-1H-pyrazolo[3,4-d]pyrimidin-4-amine (1-NM-PP1; LGC Standards, UK) was used from a 50 mM stock in DMSO. Working medium containing 5 µM 1-NM-PP1 was prepared fresh alongside a DMSO-only control (0.01% final DMSO in both cases). Medium was exchanged as described above, and z-stacks were acquired 10 minutes after exchange. For colony assays, 1-NM-PP1 (or DMSO alone) was added to ANMM agar after autoclaving and cooling to ∼60°C before pouring.

Some imaging datasets were used in multiple analyses and thus contributed to multiple figures, as follows: kymograph datasets of SsdA-mScarlet3 puncta movement in wild-type and mutant backgrounds were used in both **Fig. 2F** and **Fig. 4B**; still images of SsdA-mScarlet3 puncta in wild-type and mutant backgrounds were used in **Fig. S4C-D** and **Fig S5C**; still images of SsdA-mScarlet3 and 3xTagGFP2-RabA puncta were used in **Fig. S3F-G** (first time-point only) and **Fig. 5**.

### FRAP

For FRAP experiments, we typically bleached regions between neighboring nuclei, and pixel dwell times for photobleaching were adjusted to achieve total bleaching of a region of interest within 5–20 s. Photobleaching was performed using combined 405 nm and 561 nm laser lines.

### Image and data analysis

#### Segmentation of hyphae

Prior to quantitative analysis of fluorescent puncta, fluorescence intensity profiles, and hyphal growth rates, we generated segmentation masks of hyphae. Segmentation masks were made from maximum-intensity projections of SsdA-Scarlet3 z-stack images, using the interactive segmentation tool micro-SAM (v1.6, Archit et al., 2025) within napari (Sofroniew et al., 2026). Masks were generated using micro-SAM’s “image_series_annotator” interface with the “vit_b_lm” model (a ViT-Base SAM variant fine-tuned on light microscopy images) and corrected manually where necessary. We typically generated segmentation masks of tip-proximal regions of hyphae, excluding any hyphal branches from the mask. Segmentation masks were saved as TIFF files matching the dimensions of the original images, with pixel values of 0 for all pixels in the background and different integral pixel values for pixels within each labelled hypha, on a hypha-by-hypha basis.

#### Fluorescent puncta detection and quantification

To detect fluorescent puncta of SsdA and RabA, we used maximum-intensity z-projections and Spotiflow v0.6.4, a convolutional neural network-based spot detection tool (Dominguez Mantes et al., 2025). For spot detection in images of wild-type and mutant SsdA-mScarlet3 puncta, we generated a fine-tuned Spotiflow model, derived from the pretrained “synth_complex” base model (trained on synthetic images). Fine-tuning was performed for 30 epochs on SsdA-mScarlet3 images and their respective curator-corrected spot annotations: 20 images for model training (12 wild-type, 2 *ssdA-WAKA*, 2 *pxdAΔ*, 2 *dipAΔ*, and 2 *ssdA-6D*), and 5 images held out for model validation during training (2 wild-type, 1 *ssdA-WAKA*, 1 *dipAΔ*, 1 *ssdA-6D*). Model performance was then assessed using 4 test images that were not used for training or model validation (2 wild-type, 1 *ssdA-WAKA*, 1 *pxdAΔ*; 218 curator-corrected spots). For spot detection with the fine-tuned model, we used a confidence threshold (which can range from 0 to 1) of 0.475; this achieved F1 scores of 0.76–0.85 across genotypes (precision 0.70–0.78, recall 0.82–0.93), at a matching tolerance of 3 pixels (0.33 µm) between curator-corrected prediction and model prediction. An independent informal test of model performance was provided by analysis of SsdA-NtermΔ, which by eye displays a largely cytosolic signal, with only sparse, faint puncta (**Fig. 7A**); Spotiflow analysis of SsdA-NtermΔ showed that half of all detected SsdA-NtermΔ puncta have a fluorescent signal below that of the lowest seven percent of wild-type SsdA puncta (**Fig. S7D**). This indicates that most puncta with signals above this level are likely to be *bona fide* puncta and not artifacts of analysis.

For spot detection in images of 3xTagGFP2-RabA puncta, we used the Spotiflow “general” model and a confidence threshold of 0.6. The higher threshold for RabA puncta was possible because of the brighter, more discrete early endosome signal relative to SsdA puncta.

Positions and signals of individual fluorescent puncta were also obtained using Spotiflow, which returns subpixel xy coordinates and an integrated-intensity value for each detected spot. Spotiflow integrated-intensity values are determined by a weighting of pixel intensity values of the pixels surrounding the predicted subpixel location of the detected spot (Dominguez Mantes et al., 2025). Before plotting for figures, puncta fluorescent signals were further corrected by subtracting cytosolic background signal.

To assign detected spots to specific hyphae, we used scikit-image v0.25 in Python. We quantified puncta area number density as the number of detected spots divided by the hyphal area, measured from the segmentation mask.

#### Nearest-neighbor distance analysis for puncta colocalization

To quantify spatial relationships between SsdA-mScarlet3 and 3xTagGFP2-RabA puncta, we calculated nearest-neighbor distances using a KDTree spatial indexing algorithm (cKDTree from SciPy 1.13). The algorithm built a KDTree from the coordinates of all RabA puncta in a given hypha, then queried this tree with the coordinates of all SsdA puncta to find the closest RabA neighbor to each SsdA punctum. Distances were calculated as Euclidean distances in pixel space, then converted to microns. We classified SsdA puncta as colocalizing with RabA when the nearest-neighbor distance was ≤ 0.3 µm.

#### Skeleton generation and distance measurements

To measure hyphal length, calculate distances from fluorescent puncta to hyphal tips, and measure fluorescence intensity profiles of MobB, we extracted one-pixel-wide centerlines (skeletons) from segmented hyphae. Skeleton extraction was performed using custom Python code (Python 3.12) with scikit-image v0.25 for morphological operations and NetworkX v3.5 for graph-based analysis. For each segmented hypha, the binary mask was smoothed using a Gaussian filter (σ = 1.0) to reduce noise-induced irregularities, then skeletonized using the ‘skeletonize’ function of scikit-image. When branches appeared in skeletons (due to mask irregularities such as bulges or constrictions), we identified the longest path between all branch endpoints using Dijkstra’s algorithm and retained only skeleton pixels along this path, yielding a continuous, unbranched centerline. Skeleton extraction used identical parameters across all image analyses.

Because skeletonization erodes inward from the true hypha boundary, before using skeletons in measurements, we needed to extend each skeleton to the corresponding hyphal tip. First, hyphal tip coordinates were assigned manually with a custom napari plugin. Briefly, we overlaid maximum-intensity projections and segmentation mask outlines and placed a point at each hyphal tip. Each point was assigned to its enclosing hypha (or nearest hypha by Euclidean distance) and snapped to the closest pixel on the segmentation mask outline, yielding a tip coordinate on the segmentation mask outline. Tip coordinates were saved as JSON files for use in subsequent analysis. Second, we extended the skeleton end (closest to the tip) to the coordinates of the tip using scikit-image’s “linè function.

For calculating hyphal length and distances of puncta to hyphal tips, we converted each skeleton to a weighted graph with one node per skeleton pixel and edges connecting 8-adjacent neighbors. Edge weights were set to 1.0 for orthogonal connections and √2 for diagonal connections to correctly account for pixel geometry. Hyphal length was calculated as the sum of all edge weights multiplied by the pixel size (0.11 µm/pixel).

Distances from detected spots to tips were calculated as geodesic distances along the skeleton graph. Each spot coordinate was first projected onto the nearest skeleton node. For each spot, we calculated the distance from the projected point to the tip using NetworkX ‘shortest_path_length’.

#### MobB-mNeonGreen Hyphal Intensity Profile Analysis

To measure the spatial distribution of MobB-mNeonGreen fluorescence relative to SsdA punctate signal along the hyphal axis (**Fig. 6D**), we computationally straightened curved hyphae and extracted intensity profiles from tip to base, similarly to Vernay et al. (2012). This approach allowed consistent cross-sectional averaging along the length of curved hyphae.

To computationally straighten hyphae, we first generated skeletons from SsdA-mScarlet3 maximum-intensity projections, as described above. A smooth backbone was then created by linearly interpolating the ordered skeleton coordinates as a function of cumulative arc length. Using this backbone, we applied an elastic deformation transformation to sum-projection images of MobB-mNeonGreen.

An inverse mapping based on the local tangent and perpendicular at each backbone position was used with scikit-image’s “warp” function (bilinear interpolation) to resample the curved hypha into a rectangular image whose length matched the skeleton arc length. For each row of the straightened image (corresponding to a position along the hypha axis), we averaged background-corrected fluorescence intensities across all pixels belonging to the hypha mask, producing a one-dimensional intensity profile from tip to base. Background intensity was subtracted from all measurements.

#### Kymograph generation and analysis of movements

To quantify SsdA puncta movement, we generated kymographs from timelapse movies using the KymoResliceWide plugin (Katrukha, 2020) or KymographBuilder (Mary et al., 2016) in Fiji (Schindelin et al., 2012). For quantification of movement velocities, we first generated maximum intensity projections along the time axis, to identify regions suitable for kymographing. We then placed a 5 pixel-wide line along regions of interest and used this to generate kymographs. For quantitative comparison of movement frequency among different conditions, we instead drew 10 µm line ROIs adjacent to a nucleus, ensuring equivalent sampling regions across movies.

Kymographs were analyzed using KymoButler (Jakobs et al., 2019), a deep learning-based track detection package implemented in Wolfram Mathematica (v14.2), with standardized parameters across all analyses: detection threshold 0.1, minimum particle size 3 pixels, and minimum track duration 500 ms (10 frames when 50 ms exposure) or 480 ms (6 frames when 80 ms exposure). KymoButler calculated the mean velocity for each track as total distance traveled divided by track duration and classified the movement direction as anterograde (toward hyphal tip) or retrograde (away from hyphal tip), based on kymograph orientation.

To quantify co-localizing movements between different proteins, we generated kymographs from dual-channel timelapse videos and scored each kymograph manually, classifying each visible movement as channel 1 alone, channel 2 alone, or co-moving.

#### Manual hyphal growth rate tracking

To measure instantaneous hyphal tip growth rates before and after acute drug treatments (**Fig. 6E, S6B**), we tracked hyphal tip positions manually across time-lapse movies using the ImageJ mTrackJ plugin (Meijering et al., 2012). We followed the tracking procedure described by Athanasopoulos et al. (2021).

For each movie, we manually placed points at the hyphal tip in every timepoint, creating a trajectory of tip positions over time. Tracking data (raw x,y coordinates and timepoints) were exported as .mdf files from mTrackJ and imported into R for analysis.

We calculated tip growth rates in R as follows. Pixel coordinates were converted to microns (0.275 µm per pixel), and timepoints were converted to time (5 minutes per timepoint). For each tracked hypha, we calculated the step-by-step Euclidean distance between consecutive tip positions, then summed these incremental distances to produce a cumulative path length traveled over time. Because manual tracking introduces timepoint-to-timepoint jitter, we smoothed these cumulative distance curves using a 3-point moving average (using the “rollmean” function from the zoo R package (Zeileis and Grothendieck, 2005), centered on each timepoint). Instantaneous velocity (µm/min) was calculated as the change in smoothed distance divided by the time interval between consecutive frames.

To account for hypha-to-hypha variability in baseline growth rate, we normalized velocities to each hypha’s pre-treatment baseline velocity (mean velocity across the first 12 frames, corresponding to the first 60 minutes).

#### Relating hyphal growth rate to puncta spatial distribution

To measure hyphal growth rates and correlate them with puncta localization patterns (**Fig. 5**), we acquired two-channel fluorescence images (SsdA-mScarlet3 and 3xTagGFP2-RabA) at two timepoints (T0 and T1) separated by 25 minutes. We used maximum-intensity projections, hypha segmentation masks and manually selected hyphal tips as described above.

To track individual hyphae across the 25-minute interval, we matched T1 segmentation masks to T0 segmentation masks using the Hungarian algorithm (Kuhn, 1955), which finds optimal one-to-one assignments by identifying maximal spatial overlap between segmentation masks. We quantified overlap as T0 coverage (intersection area / T0 area), which is appropriate for growing hyphae, for which the T1 mask is expected to be larger than the T0 mask. Hypha matching was implemented using SciPy’s “linear_sum_assignment” function (SciPy 1.13). For each matched hypha pair, we extracted skeletons as described above. To calculate the extent of growth during the 25-minute interval, we measured the geodesic distance along the T1 skeleton from the T1 tip position to the point (on the T1 skeleton) that was nearest to the T0 tip position. Growth rate (µm/min) was calculated as geodesic distance in µm divided by 25 min.

#### Calculating length of SsdA depletion zone

To quantify SsdA depletion zones at hyphal tips, we first determined the position and signal of all puncta (as described above) within 50 µm of the tip of a given hypha. We defined the sum of the signals from these puncta as the “total punctate signal” (i.e. within 50 µm of the tip) of a given hypha. We then sorted the puncta based on their geodesic distance along a skeleton starting at the hyphal tip, and we calculated the cumulative sum of punctum signals as a function of distance from the tip. We defined the length of the depletion zone as the distance along the skeleton (from the tip) at which the cumulative puncta signal first reached 5% of the total punctate signal.

Because puncta are present at discrete distances along the skeleton, we determined the distance corresponding to 5% by linear interpolation between the two puncta on either side of 5%. We prefer this approach over simply measuring the distance of the first punctum closest to the hyphal tip, which would be particularly sensitive to faint, outlier or false-positive puncta.

We used the exact same approach for RabA puncta.

Because photobleaching reduced punctum counts at T1 (particularly for SsdA-mScarlet3), when we correlated depletion zone length with hyphal growth rate, we first calculated the depletion zone length separately for each timepoint and then used the mean of the two timepoints as a single per-hypha value (**Fig. 5E**).

### Bioinformatic analysis of Ssd1 binding motif

The annotated transcriptome of *A. nidulans* was obtained from FungiDB version 64 (Basenko et al., 2018). For genes with multiple annotated 5′ untranslated region isoforms, we selected the longest UTR for analysis. We counted occurrences of the Ssd1 consensus binding motif ’CNYTCNYT’ (equivalent to ’CNYUCNYU’ in RNA annotation; Bayne et al., 2022) in annotated 5′ UTRs using the Biostrings package v.2.74.1 in R, using similar code as in Bayne et al. (2022). Genes with four or more Ssd1 binding sites were compiled into a list and submitted to FungiDB for Gene Ontology “Cellular Component” enrichment analysis (Basenko et al., 2018). Terms annotated as ’obsolete’ were excluded from the results.

### Protein structure prediction and visualization

Protein structure predictions were generated with AlphaFold3 using the AlphaFold Server (Abramson et al., 2024) and downloaded for visualization in PyMOL (Schrödinger, LLC, 2015). Predicted aligned error (PAE) matrices were visualized using PAE Viewer (Elfmann and Stülke, 2023).

### Data visualization and statistics

All analyses were performed in R (v4.5) using ggplot2 (v4.0; Wickham, 2016) for visualization. Each biological replicate comprised an independent experiment using hyphae derived from a separate colony streak. Graphs were typically plotted as ‘Superplots’, following Lord et al. (2020). Statistical comparisons were made within each biological replicate; significance was reported only when all replicates independently passed the same threshold (p < 0.001, p < 0.01, p < 0.05). Correlation of growth rate with puncta depletion zone length and puncta number (**Fig. 5**) was analyzed using pooled data from the three biological replicates. Biological replicate numbers and statistical tests performed are reported in figure legends.

## Supporting information

Combined supplemental information

## Software and data availability

Data and code for reproduction of analysis have been deposited in Zenodo (doi: 10.5281/zenodo.19401665). Portions of code were generated with assistance from Claude (Anthropic).

## Acknowledgments

We thank Dhanya Cheerambathur for use of the dual-camera Nikon Ti2 spinning disk microscope, Xin Xiang and Sam Reck-Peterson for strains, Berl Oakley for valuable advice on *A. nidulans* techniques, and members of the Sawin and Wallace labs for advice on the project as it has developed. This work was supported by funding for the Wellcome Discovery Research Platform for Hidden Cell Biology. We gratefully acknowledge support from the light microscopy core technologists Dave Kelly and Toni McHugh for help and advice on microscopy equipment and software. For the purpose of open access, the authors have applied a Creative Commons Attribution (CC BY) licence to any Author Accepted Manuscript version arising from this submission.

