## Supplementary material for "Endosomal hitchhiking and NDR kinase signaling coordinate SsdA-mRNP localization": Combined supplemental information

### Video legends

**Video S1. SsdA forms motile cytoplasmic puncta.**
Single-z-section continuous-acquisition movies of SsdA tagged with mScarlet3 (top) and mNeonGreen (bottom). The SsdA-mScarlet3 hypha is the same as shown in Fig. 1D. With either fluorescent protein tag, SsdA forms motile cytoplasmic puncta. Scale bar = 5 μm.

**Video S2. FRAP allows visualization of motile FabM puncta.**
Single-z-section continuous-acquisition movies of poly(A)-binding protein FabM-mScarlet3 in a FRAP experiment. Before photobleaching, few to no motile FabM puncta are observed. After photobleaching, FabM puncta are readily observed moving into dark (bleached) area. The same hypha is shown in Fig. S2C. Scale bar = 5 μm

**Video S3. SsdA puncta move on microtubules.**
Single-z-section simultaneous continuous-acquisition movies of hypha expressing SsdA-mScarlet3 (magenta) and GFP-TubA (green). Arrowheads indicate examples of SsdA puncta moving on microtubules. The same hypha is shown in Fig. 3A and Fig. S3A. Scale bar = 5 μm.

**Video S4. SsdA puncta move with early endosomes.**
Single-z-section simultaneous continuous-acquisition movies of hypha expressing SsdA-mScarlet3 (magenta) and 3xTagGFP2-RabA (green; early endosome marker). Arrowheads indicate examples of colocalizing movements. The same hypha is shown in Fig. 3D. The SsdA-mScarlet3 channel was bleach-corrected for easier visualization. Scale bar = 5 μm.

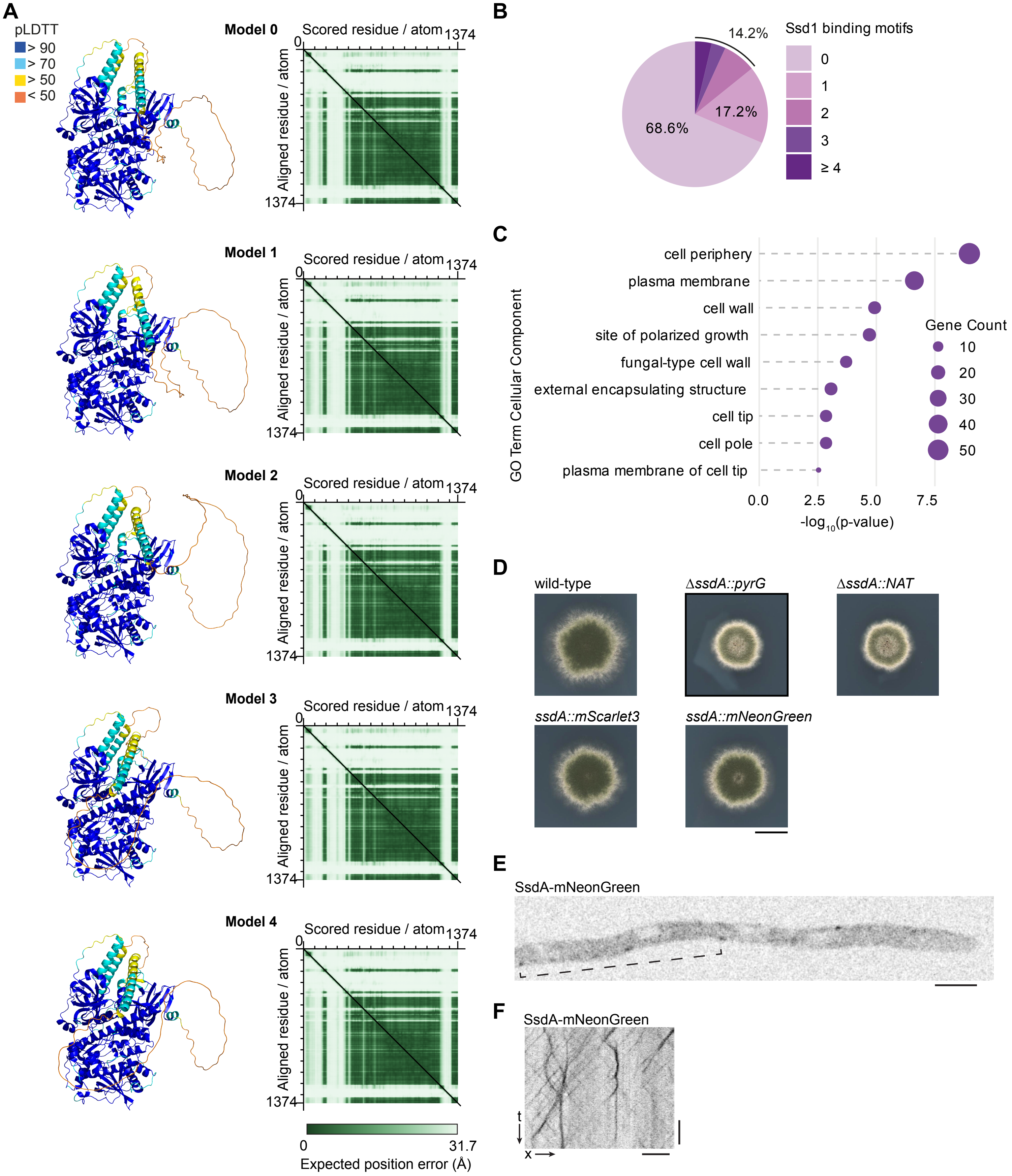

**Figure S1. Additonal data related to Figure 1. (A)** AlphaFold3 structural predictions of SsdA, showing five model variants (left) with corresponding predicted aligned error (PAE) plots (right). Structural predictions are colored by pLDDT (predicted local distance difference test) score. PAE matrices are colored by expected position error between residue pairs. **(B)** Distribution of numbers of Ssd1 consensus binding motifs "CNYUCNYU" across annotated *A. nidulans* 5' untranslated regions (5' UTRs) (n = 7072 genes). **(C)** Gene Ontology (GO) "Cellular Component" enrichment analysis for *A. nidulans* genes containing ≥ 4 Ssd1 consensus binding motifs in their 5' UTRs (n = 261 genes. Y-axis includes all statistically significant identified GO terms. X-axis shows Bonferroni-adjusted p-value. **(D)** Colony phenotypes of wild-type *A.*

*nidulans*, two independent *ssdA* deletion strains (using either *pyrG* or *NAT* as selectable markers), and two different C-terminal fluorescent-tagged alleles of *ssdA* (tagged at endogenous locus). Colonies were grown on minimal medium for 3 days at 32°C. Both tagged alleles of *ssdA* resemble wild-type, unlike either *ssdA* deletion strain. Scale bar = 1 cm. **(E, F)** Zero timepoint image from single-z-section movie of hypha expressing SsdA-mNeonGreen from endogenous locus (E), with accompanying kymograph (F). Bracket in (E) indicates region kymographed in (F). Scale bars = 5 μm, 2 s.

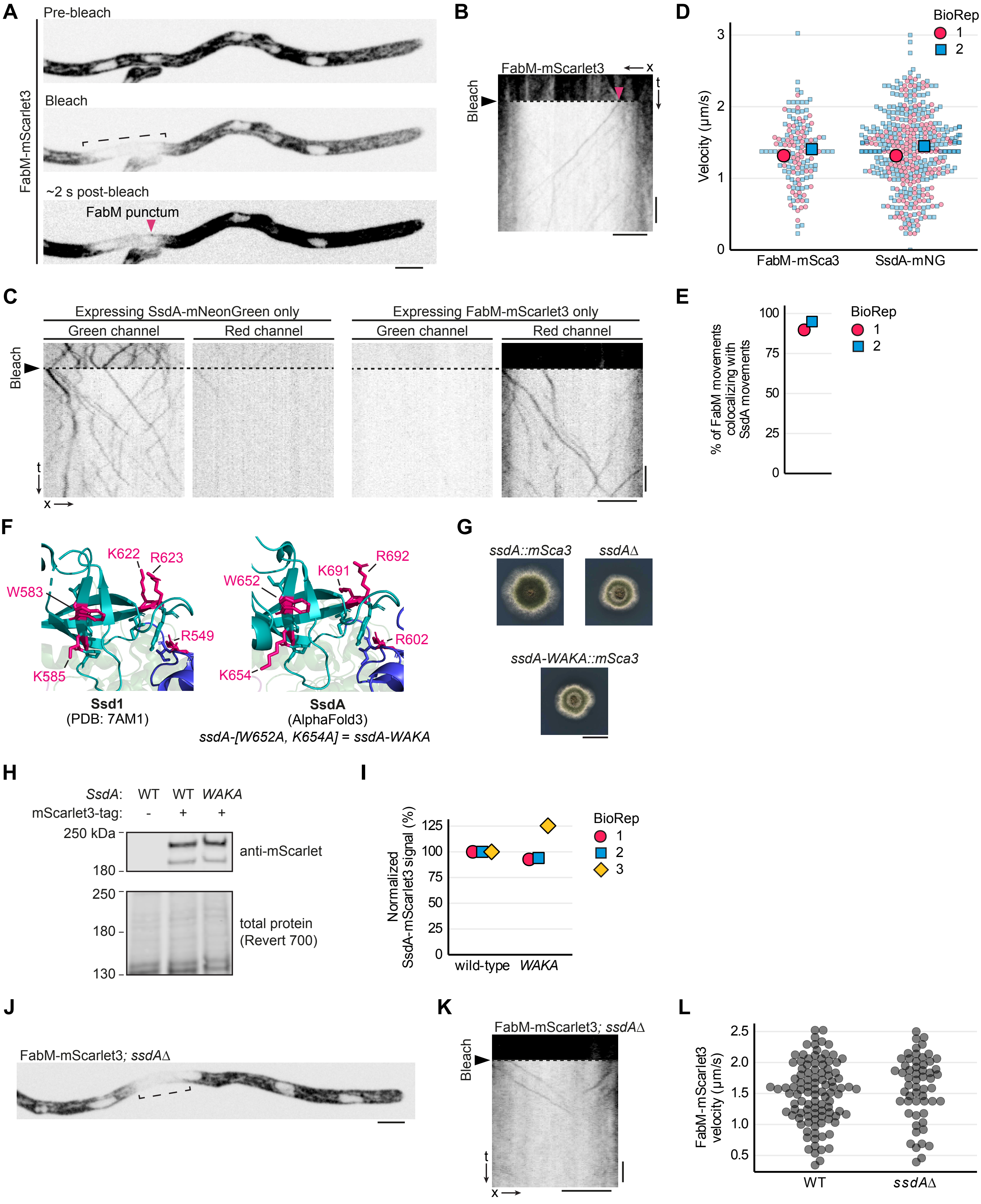

**Figure S2. Additonal data related to Figure 2. (A, B)** Timepoints from single-z-section movie of hypha expressing poly(A)-binding protein FabM-mScarlet3 from endogenous locus before (pre-bleach), immediately after (bleach), and ~2 seconds after photobleaching (post-bleach) (A), with accompanying kymograph (B). Bracket in (A) indicates bleached region, which is kymographed in (B). Magenta arrowhead indicates moving FabM-mScarlet3 punctum. Post-bleach timepoint in (A) is contrast-enhanced compared to pre-bleach and bleach timepoints. **(C)** Kymographs from dual-color imaging experiments as in Fig. 2A, but with strains expressing either only SsdA-mNeonGreen (left) or only FabM-mScarlet3 (right), to test for fluorescence bleed-through. SsdA-mNeonGreen puncta exhibit very low bleed-through into the red

channel; any bleed-through that is observed is restricted to the very brightest SsdA-mNeonGreen puncta. FabM-mScarlet3 puncta do not exhibit any observable bleed-through. These imaging conditions thus allow colocalization measurements with negligible false positives. **(D)** Velocity distributions of SsdA-mNeonGreen and FabM-mScarlet3 puncta, quantified from kymographs generated from photobleaching movies of hyphae expressing both SsdA-mNeonGreen and FabMmScarlet3 as in Fig. 2A, B. Small symbols indicate individual puncta movements; large symbols indicate means of independent biological replicates (BioReps). Different shapes/colors indicate distinct BioReps. Numbers of movements scored: 64, 79 for FabM-mScarlet3 and 266, 246 for SsdA-mNeonGreen in BioReps 1, 2 respectively. Velocities of FabM-mScarlet3 and SsdA-mNeonGreen puncta were statistically indistinguishable (two-tailed Welch's unpaired t-test, $p > 0.05$ for each BioRep), consistent with their co-transport. **(E)** Percentage of FabM-mScarlet3 movements observed to colocalize with SsdA-mNeonGreen movements, quantified from kymographs as in (D) (see Fig. 2B). Number of FabM movements scored = 78, 99 in BioReps 1, 2 respectively. Nearly all detectable moving FabM-mScarlet3 puncta contain detectable SsdA-mNeonGreen. **(F)** Residues shown to be involved in RNA binding (colored in pink) in *S. cerevisiae* Ssd1 (Bayne et al., 2022), with corresponding conserved residues in *A. nidulans* SsdA. Ssd1's K585 has a few atoms missing from its side chain, as they were not resolved in the crystal structure. Remainder of structures are colored as in Fig. 1B. We describe the *ssdA* double mutant W652A, K654A, corresponding to the *S. cerevisiae* Ssd1 CSD-TOP mutant W583A, K585A (Bayne et al., 2022), as *ssdA-WAKA*. **(G)** Colonies of *ssdA::mScarlet3* (wild-type), *ssdA-WAKA::mScarlet3*, and *ssdAΔ* (*ssdAΔ::NAT*) strains, grown from spores on minimal medium for 3 days at 32°C. The *ssdA-WAKA::mScarlet3* phenotype resembles *ssdAΔ*. **(H, I)** Representative anti-mScarlet western blot of TCA extracts from untagged wild-type strain and mScarlet3-tagged wild-type and *ssdA-WAKA* strains (H), with quantitation from three independent BioReps (I). Signal in (H) is normalized to total protein (Revert700 stain) and further normalized to wild-type signal within each BioRep. Different shapes/colors indicate distinct BioReps. **(J, K)** Photobleaching of FabM-mScarlet3 in an *ssdA* deletion strain (*ssdAΔ::NAT*). Timepoint from single-z-section movie immediately after bleaching (J), with accompanying kymograph (K). Bracket in (J) indicates bleached region, which is kymographed in (K). FabM-mScarlet3 puncta movement does not require *ssdA*. **(L)** Velocity distributions of FabM-mScarlet3 puncta in wild-type and *ssdAΔ* backgrounds. Symbols indicate individual puncta movements. FabM puncta velocities are indistinguishable between wild-type and *ssdAΔ* backgrounds (two-tailed Welch's unpaired t-test, $p > 0.05$). Number of movements were 101 movements for wild-type, and 56 for *ssdAΔ*.
Scale bars = 5 µm, 2 s for micrographs and kymographs, 1 cm for colony pictures.

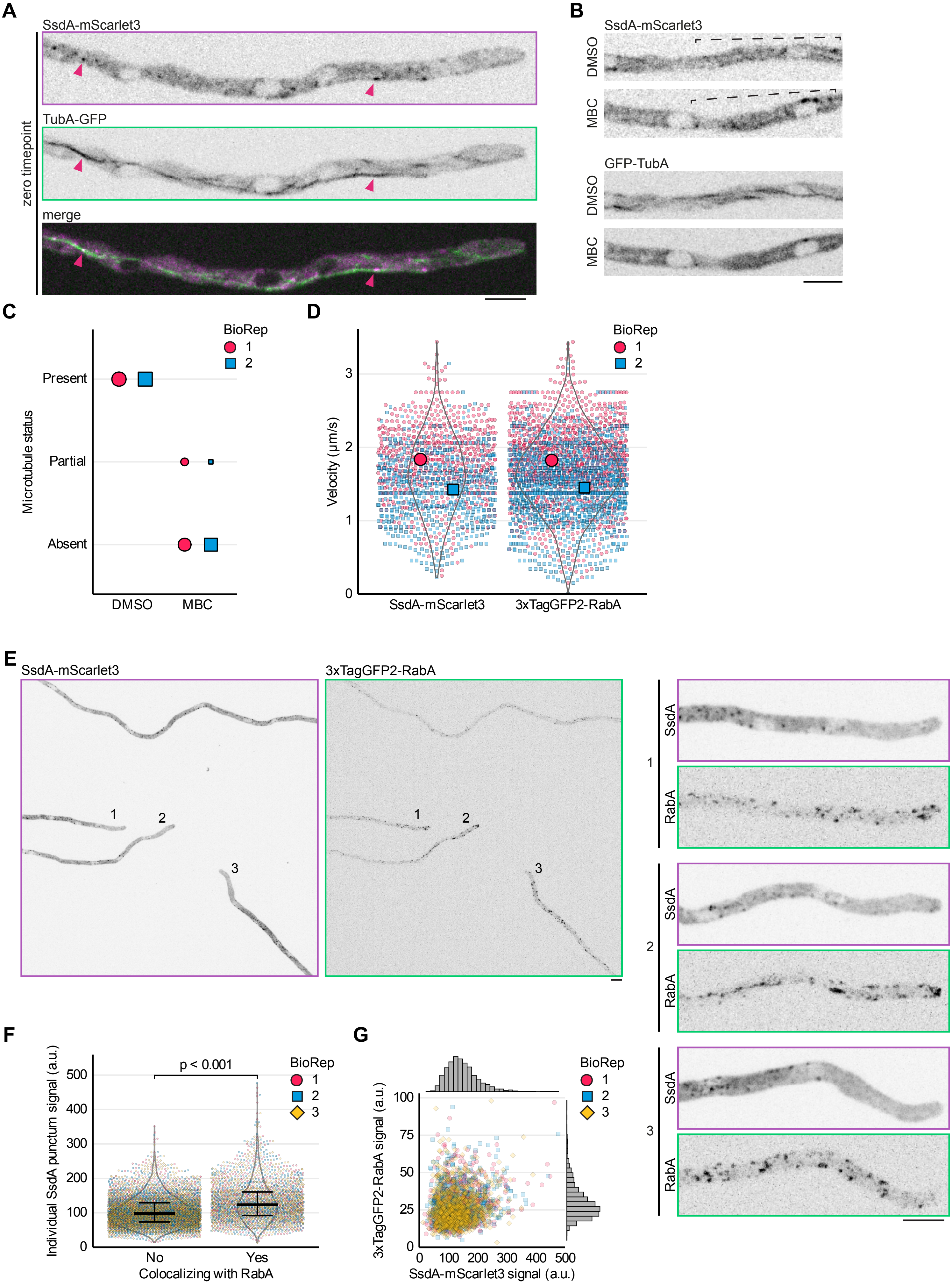

**Figure S3. Additonal data related to Figure 3.** Figure legend on next page.

**Figure S3. (A)** Zero timepoint from single-z-section movie used to make time projection shown in Fig. 3A. Arrowheads indicate examples of SsdA puncta on microtubules. **(B)** Zero timepoints from single-z-section movies of hyphae expressing SsdA-mScarlet3 and TubA-GFP, 10 minutes after exchange to fresh medium containing either 25 μg/mL MBC or vehicle (DMSO) alone. Brackets indicate kymographed regions of SsdA-mScarlet3, shown in Fig. 3B. **(C)** Microtubule status in hyphae expressing SsdA-mScarlet3 and TubA-GFP, 10 minutes after exchange to fresh medium containing either 25 μg/mL MBC or DMSO alone. Area of symbols indicate relative proportion of imaged fields where microtubules were completely depolymerized (absent), where at least one hypha showed at least one intact microtubule (partially present), or where all hyphae in the field had intact microtubules (present). Different shapes/colors indicate distinct biological replicates (BioReps). Number of fields imaged: 13, 17 for DMSO, and 13, 16 for MBC in BioRep1, 2 respectively.
**(D)** Velocity distributions of SsdA-mScarlet3 and early endosome marker 3xTagGFP2-RabA puncta, quantified from 5 s kymographs of the type shown in Fig. 3D. Small symbols indicate individual puncta movements; large symbols indicate means of BioReps. Different shapes/colors indicate distinct BioReps. Numbers of movements scored: 600, 592 for SsdA-mScarlet3 and 1020, 1120 for 3xTagGFP2-RabA in BioReps 1, 2 respectively. Velocities of SsdA-mScarlet3 and 3xTagGFP2-RabA puncta were statistically indistinguishable (unpaired two-tailed Welch's t-test, $p > 0.05$ for each BioRep), consistent with their co-transport. **(E)** Representative field of view of hyphae expressing SsdA-mScarlet3 and 3xTagGFP2-RabA. Maximum-intensity z-projection of simultaneous dual-channel acquisition. Zoomed images of numbered hyphae are shown at right. **(F)** Comparison of SsdA-mScarlet3 puncta signal between puncta not colocalizing and puncta colocalizing with 3xTagGFP2-RabA. Colocalization was defined as centroid distance between SsdA and RabA puncta <0.3 μm. Lines and error bars represent median and interquartile range. Area of violin plots is proportional to the total number of puncta in each of the two categories. Different shapes/colors indicate distinct BioReps. Significance brackets indicate thresholds met in all BioReps (Mann–Whitney *U*, two-tailed, per BioRep). Number of spots were 1602, 2593, and 3362 for BioReps 1 to 3. **(G)** Comparison of fluorescence signal between colocalizing 3xTagGFP2-RabA and SsdA-mScarlet3 puncta. Distribution of puncta signals are plotted as marginal histograms on corresponding axes. Colocalization was defined as in (F). No strong correlation between fluorescence signals is observed (Pearson's $r = 0.19$, $p < 0.001$). Number of colocalizing pairs were 696, 847, and 1107, for BioReps 1 to 3.
Imaging datasets used to generate panels F and G were also used for analyses of SsdA and RabA puncta shown in Fig. 5.
Scale bars = 5 μm

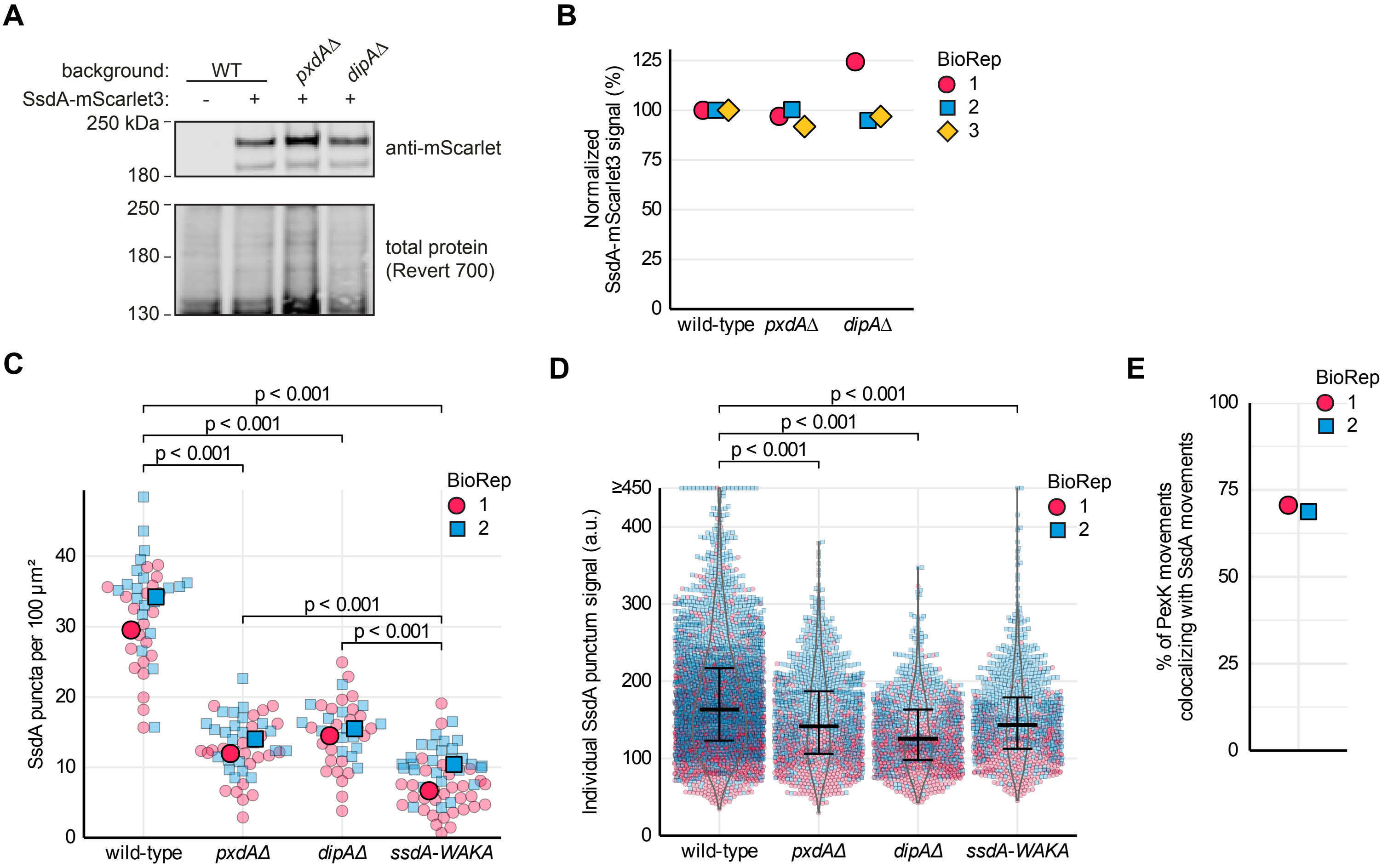

**Figure S4. Additonal data related to Figure 4. (A, B)** Representative anti-mScarlet western blot of TCA extracts from untagged wild-type strain and mScarlet3-tagged wild-type, *pxdAΔ* and *dipAΔ* strains (A), with quantitation from three independent biological replicates (BioReps) (B). Signal in (B) is normalized to total protein (Revert700 stain) and further normalized to wild-type signal within each BioRep. Different shapes/ colors indicate distinct BioReps.

**(C)** Area number density of SsdA-mScarlet3 puncta in wild-type, hitchhiking-adaptor deletion mutants *pxdAΔ* and *dipAΔ,* and predicted RNA-binding mutant *ssdA-WAKA*. Puncta were scored within regions of hyphae ranging between ~50 and ~400 µm$^2$ and values were then scaled to units of puncta per 100 µm$^2$. Small symbols indicate individual hyphae; large symbols indicate means of BioReps. Different shapes/ colors indicate distinct BioReps. Brackets indicate significant two-tailed unpaired Welch's t-tests. Number of hyphae: wild-type = 21, 21; *pxdAΔ* = 23, 24; *dipAΔ* = 25, 15; *ssdA-WAKA* = 31, 27 for BioRep 1, 2 respectively.

**(D)** SsdA-mScarlet3 puncta signal in wild-type, hitchhiking adaptor deletions, and non-RNA-binding mutant *SsdA-WAKA*. Points represent individual background-subtracted punctum intensity. Lines and error bars indicate median and interquartile range. Statistical comparisons were performed using two-tailed Welch's unpaired t-tests separately for each BioRep. Analysis used the same imaging datasets as in (C). Number of puncta: wild-type = 1628, 1964; *pxdAΔ* = 556, 795; *dipAΔ* = 697, 556; *ssdA-WAKA* = 427, 691 for BioRep 1, 2 respectively. Compared to wild-type, all the mutants contain fewer bright SsdA puncta. **(E)** Percentage of PexK-GFP movements observed to colocalize with SsdA-mScarlet3 movements, quantified from kymographs of the types shown in Fig. 4D. Number of PexK movements scored: 17, 49 for BioReps 1 and 2, respectively.

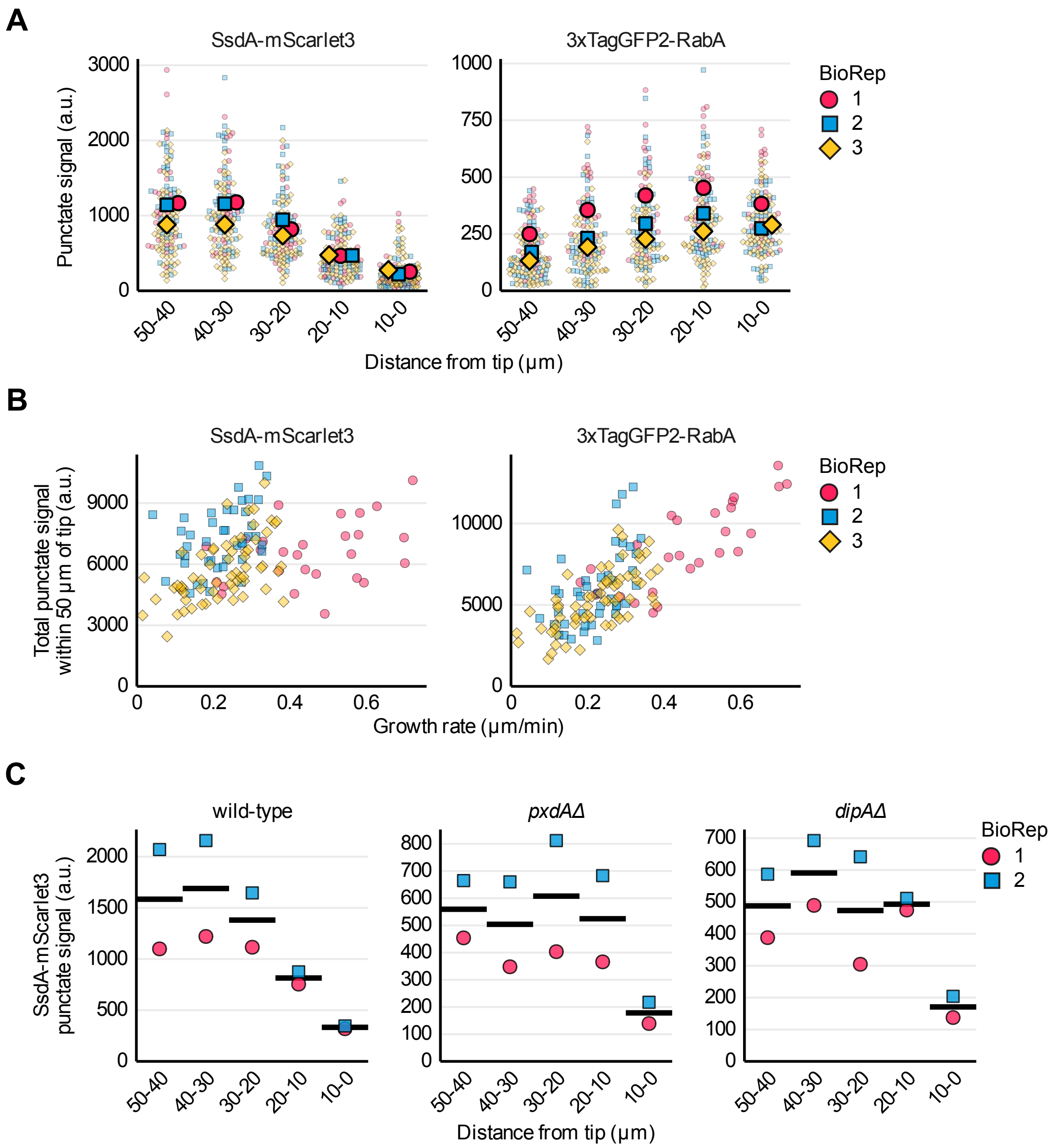

**Figure S5. Additonal data related to Figure 5. (A)** SsdA-mScarlet3 and 3xTagGFP2-RabA punctate signal (sum of individual puncta signal) in 10 µm-long segments within the 50-µm region of hyphae closest to hyphal tips. Small symbols indicate individual hyphae; large symbols indicate means of biological replicates (BioReps). Different shapes/colors indicate distinct BioReps. Analysis used the same imaging datasets as in Fig. 5. **(B)** Total signal of SsdA-mScarlet3 or 3xTagGFP2RabA puncta in the 50-µm region closest to hyphal tips, plotted against hyphal growth rate. Symbols indicate individual hyphae. Different shapes/colors indicate distinct BioReps. Analysis used the same imaging datasets as in Fig. 5. **(C)** Mean total SsdA-mScarlet3 punctate signal in 10 µm-long segments within the 50-µm region closest to hyphal tips, in wild-type and hitchhiking-adaptor deletion mutants *pxdAΔ* and *dipAΔ*. Symbols indicate means of hyphae within each BioRep; bars indicate means of BioReps. Values are arbitrary units (a.u.). Analysis used the same imaging datasets as in Fig. S4C, D.

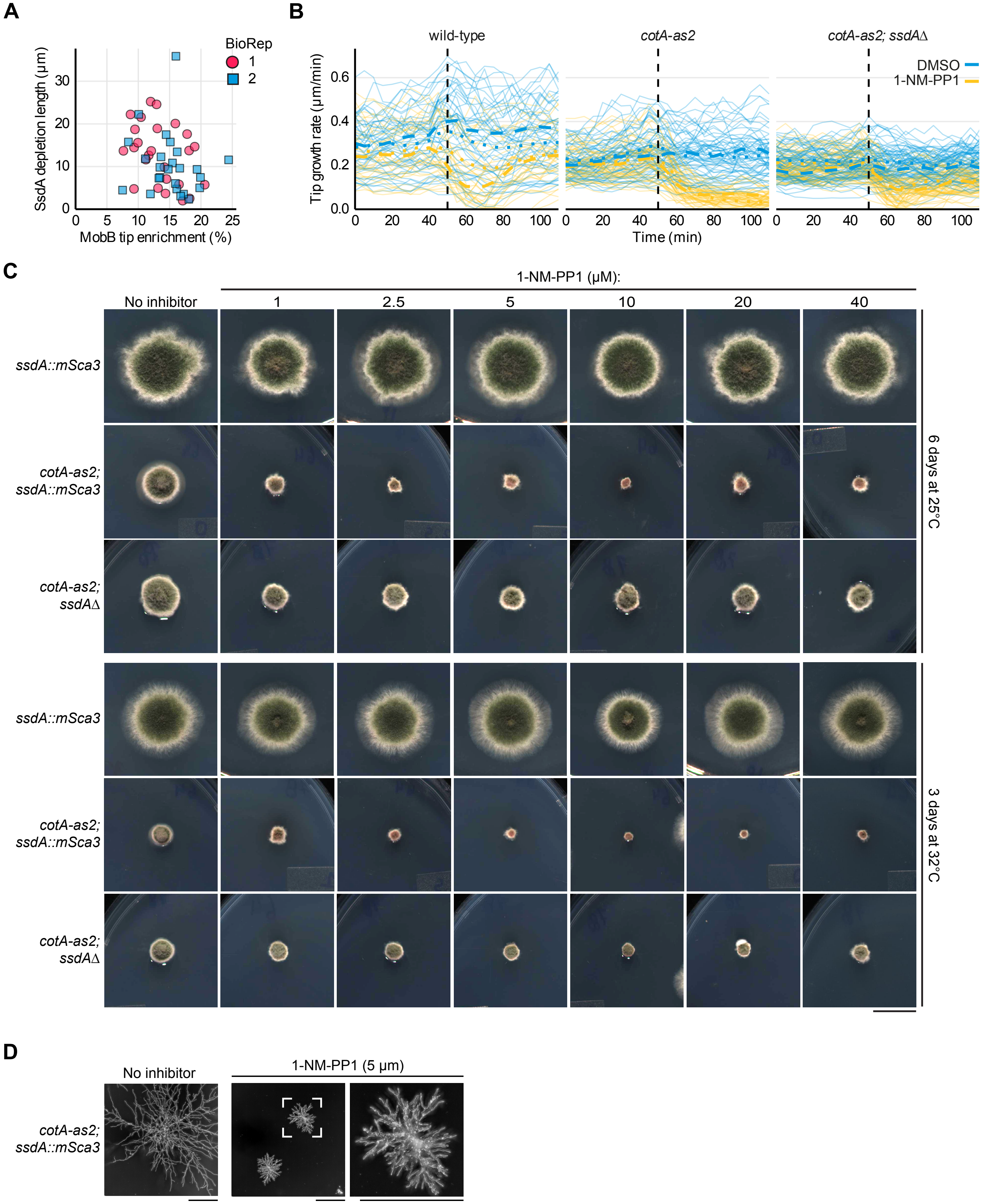

**Figure S6. Additonal data related to Figure 6. (A)** Comparison of SsdA puncta depletion and MobB enrichment near hyphal tips. SsdA tip-proximal depletion zone length was calculated as in Fig. 5D (distance from tip to reach 5% of total punctate signal within tip-proximal 50-µm region). MobB tip enrichment was measured as the percentage of total MobB-mNeonGreen signal (within the tip-proximal 50 µm) that is present within the tip-proximal 5 µm. Different shapes/colors indicate distinct biological replicates (BioReps). Analysis used the same imaging datasets as in Fig. 6D. No positive relationship between SsdA depletion zone length and MobB tip enrichment is observed. **(B)** Changes in hyphal tip growth rates upon acute inhibition of CotA. Data are identical to Fig. 6E but show absolute values rather than normalization to pre-treatment growth rates. Growth rates are shown for wild-type, *cotA-as2* (analog-sensitive CotA), and *cotA-as2; ssdAΔ* strains before and after growth-medium exchange to fresh medium containing either vehicle (DMSO) or analog-sensitive kinase inhibitor 1-NM-PP1 (5 μM). Vertical line indicates time of medium exchange. Plotted traces are three-point moving averages of raw measurements. Blue and yellow lines indicate DMSO- and 1-NM-PP1-treated hyphae, respectively. Thin lines indicate individual hyphae; thick dotted and dashed lines indicate means within each BioRep. Strains *cotA-as2* and *cotA-as2; ssdAΔ* mutations appear to grow more slowly than wild-type. **(C)** Deletion of *ssdA* partially rescues the growth and morphological defects of inhibited *cotA-as2* strains. Colony phenotypes for the indicated genotypes, grown on minimal medium containing different concentrations of 1-NM-PP1 for either 6 days at 25°C or 3 days at 32°C. The single-mutant *cotA-as2* strain exhibits dose-dependent growth inhibition and morphological defects in the presence of 1 μM or higher 1-NM-PP1, with effects plateauing beyond 5 μM 1-NM-PP1. These defects are partially rescued by further deletion of *ssdA*. In the absence of 1-NM-PP1, *cotA-as2* strains also exhibit some temperature sensitivity. Scale bar = 1 cm. **(D)** Inhibition of CotA kinase activity leads to hyperbranched colonies. Single-mutant *cotA-as2* colonies grown for 2 days at 25°C in absence of inhibitor or in presence of 5 μM 1-NM-PP1. Micrographs are minimum intensity projections of differential interference contrast microscopy images. For 5 μM 1-NM-PP1, expanded view of a colony is shown at right. Scale bar = 200 μm.

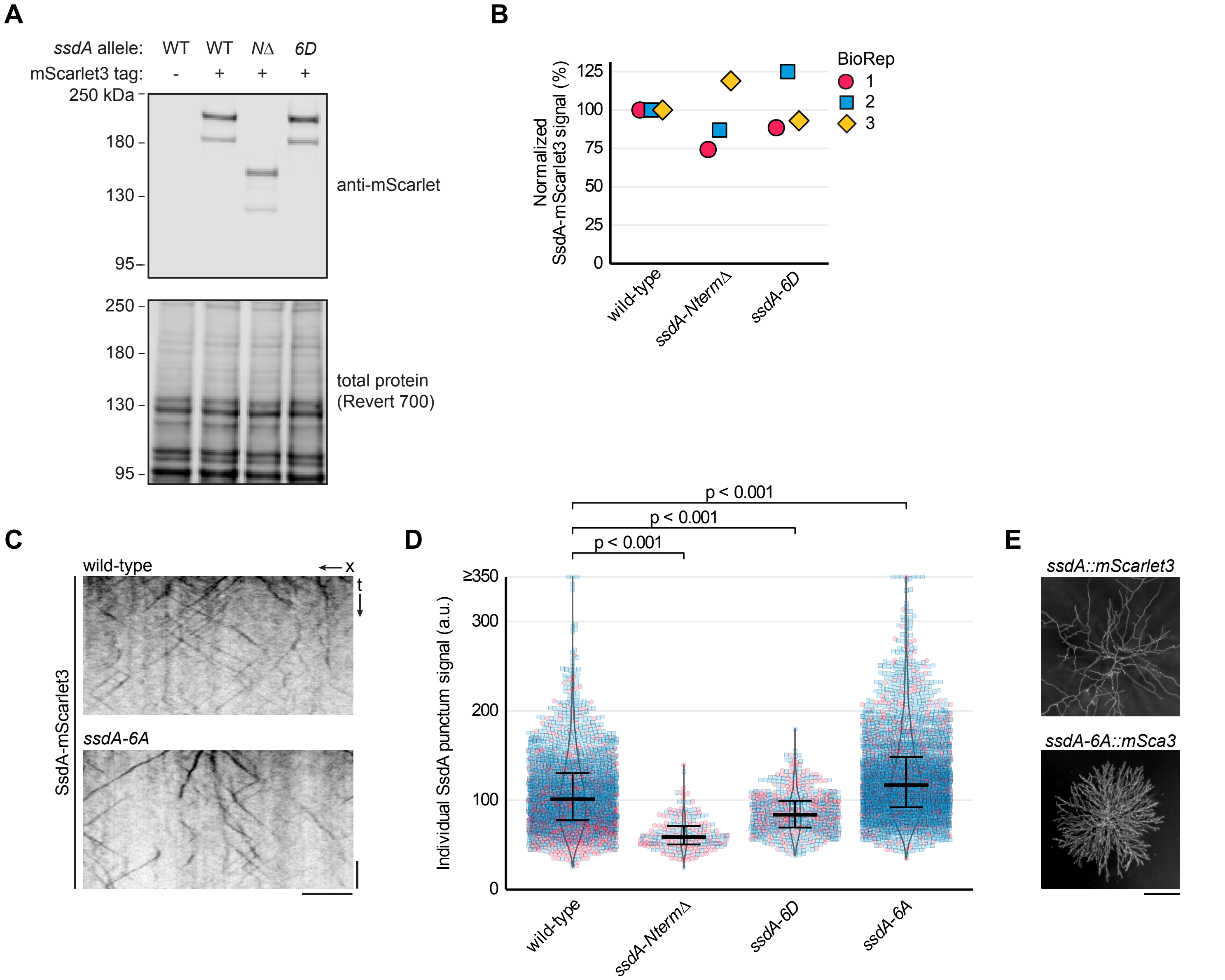

**Figure S7. Additonal data related to Figure 7. (A, B)** Representative anti-mScarlet western blot of TCA extracts from untagged wild-type strain and mScarlet3-tagged wild-type, ssdA-*NtermΔ* (*ΔN*;aa2-429Δ truncation) and *ssdA-6D* (*6D*; serine residues at NDR kinase consensus sites mutated to aspartic acid) strains (A), with quantitation from three independent biological replicate experiments (B). Signal in (B) is normalized to total protein (Revert700 stain) and then to wild-type signal within each BioRep. Different shapes/colors indicate distinct BioReps. **(C)** Representative kymographs from single-z-section continuous acquisition movies of hyphae expressing wild-type or *ssdA-6A* (serine residues at NDR kinase consensus sites mutated to alanine) SsdA-mScarlet3. SsdA-6A puncta exhibit motility comparable to wild-type. Scale bars = 5 µm, 2 s. **(D)** SsdA-mScarlet3 puncta signal in wild-type hyphae and the indicated *ssdA* N-terminal mutants. Small symbols represent individual background-subtracted puncta signals; large symbols indicate means of BioReps. Different shapes/colors indicate distinct BioReps. Values are arbitrary units (a.u.). The same dataset was analyzed as in Fig. 7C. Significance brackets indicate thresholds met in both BioReps (unpaired Welch's t-test, two-tailed, per BioRep). SsdA-mScarlet3 puncta in *ssdA-6A* are ~20% brighter than wild-type. Number of puncta: wild-type = 1961, 1604; *ssdA-NtermΔ* = 137, 106; *ssdA-6D* = 307, 388; *ssdA-6A* = 933, 2446; for BioRep 1, 2 respectively.
**(E)** Nonphosphorylatable mutation of serine residues at NDR kinase consensus sites leads to hyperbranched colonies. Early-stage colonies of *ssdA::mScarlet3* (wild-type) and *ssdA-6A::mScarlet3* strains grown for two days at 25°C. Micrographs are minimum intensity projections of differential interference contrast microscopy images. Early-stage *ssdA-6A::mScarlet3* colonies resemble CotA-kinase-inhibited colonies (Fig. S6D). Scale bar = 200 µm.

**Table S1.** Strains used in this study.

| Identifier | Genotype<br>(sc = sterigmatocystin cluster(AN7804-7825)) | Origin | Appears in figures: |
|---|---|---|---|
| Parental / reference strains | | | |
| XX872 | yA2; pyrG89; pantoB100; GFP::tubA | Xin Xiang (Qiu et al., 2022) | - |
| RPA465 | yA1, pabaA1, pyrG89; RabA(p)::3xTagGFP2::RabA5A:: AFpyrG; pyroA4, nkuA::argB | Samara Reck-Peterson (Egan et al., 2012) | - |
| RPA179 | yA1; pyrG89; pyroA4, nkuA::argB, scΔ; trpC::[pexK::GFP::pyroA] | Samara Reck-Peterson (Salogiannis et al. 2016) | - |
| aDM017 | pyrG89; pyroA4, nkuA::argB, scΔ | Previous work (Modaffari et al., 2024) | S1F, S2H, S4A, S7A |
| Core SsdA reporter strains | | | |
| aDM019 | pyrG89; pyroA4, nkuA::argB, scΔ; ssdA::mScarlet3 | Previous work (Modaffari et al., 2024) | 1(D-F), 2(C-F), 4(A, B), 6(E-I), 7(A-D), S1F, S2(G-I), S4(A-D), S5C, S6(B-D), 7(A-D) |
| aDM088 | pyrG89; pyroA4, nkuA::argB, scΔ; ssdA::mNeonGreen | CRISPR edit of aDM017 using pDM044 gRNA; repair template: pDM155 (oDM664/665) | S1(D-F), S2C |
| *ssdA* loss or functional mutants | | | |
| aDM039 | pyrG89; pyroA4, nkuA::argB, scΔ; ssdAΔ::pyrG | Homologous recombination of aDM017; repair template: pDM106 (oDM439-444) | S1F |
| aDM059 | pyrG89; pyroA4, nkuA::argB, scΔ; ssdAΔ::NAT | Homologous recombination of aDM017; repair template: pDM145 (oDM439-444) | S1F, 7B |
| aDM080 | pyrG89; pyroA4, nkuA::argB, scΔ; ssdA-NtermΔ(aa2-429Δ)::mScarlet3 | Homologous recombination of aDM039; repair template: pDM159 (oDM439-444) | 7(A-C), S7 |
| aDM073 | pyrG89; pyroA4, nkuA::argB, scΔ; ssdA-WAKA(W652A, K654A)::mScarlet3 | CRISPR edit of aDM019 using pDM128 gRNA; repair template: annealed oDM635/636 | 2(C-F), S2(G-I), S4(C, D) |
| aDM086 | pyrG89; pyroA4, nkuA::argB, scΔ; ssdA-6D(S26D, S126D, S183D, S255D, S300D, S339D)::mScarlet3 | Homologous recombination of aDM039; repair template: pDM161 (oDM439-444) | 7(A-C), S7 |
| aDM089 | pyrG89; pyroA4, nkuA::argB, scΔ; ssdA-6A(S26A, S126A, S183A, S255A, S300A, S339A)::mScarlet3 | Homologous recombination of aDM039; repair template: pDM160 (oDM439-444) | 7(A-D), S7 |
| Pathway-perturbation, compound and other strains | | | |
| aDM062 | pyrG89; pyroA4, nkuA::argB, scΔ; cotA-as2(M281A)::pyrG::cotA; ssdA::mScarlet3 | Ends-in homologous recombination of aDM019; repair template: pDM116 (oDM538/539) | - |
| aDM064 | pyrG89; pyroA4, nkuA::argB, scΔ; cotA-as2(M281A); ssdA::mScarlet3 | Selection for pyrG excision in self-cross of aDM062 | 6(G-I), S6(B-D) |
| aDM078 | pyrG89; pyroA4, nkuA::argB, scΔ; cotA-as2(M281A); ssdAΔ::pyrG | Homologous recombination of aDM064; repair template: pDM106 (oDM439-444) | 6(E, F), S6(B, C) |
| aDM094 | yA1, pabaA1, pyrG89; RabA(p)::3xTagGFP2::RabA5A:: AFpyrG; pyroA4, nkuA::argB; ssdA::mScarlet3 | CRISPR edit from RPA465 using pDM165 gRNA; repair template: pDM042 (oDM185/186) | 3(D, E), 5(B, C, E, F), S3(D-G), S4, S5(A-B) |
| aDM070 | pyrG89; pyroA4, nkuA::argB, scΔ; ssdA::mScarlet3, mobB::mNeonGreen | CRISPR edit of aDM019 using pDM137 gRNA; repair template: pDM052 (oDM616/617) | 6(B-D), S6A |
| aDM058 | pyrG89; pyroA4, nkuA::argB, scΔ; trpC::[pexK::GFP::pyroA], ssdA::mScarlet3 | Cross between aDM019 and RPA179 | 4(C, D), S4E |
| aDM049 | pyrG89; pyroA4, nkuA::argB, scΔ; pxdAΔ::pyrG, ssdA::mScarlet3 | Homologous recombination of aDM019; repair template: pDM112 (oDM474-478) | 4(A, B), S4(A-D), S5C |
| aDM056 | pyrG89; pyroA4, nkuA::argB, dipAΔ::pyrG, scΔ; ssdA::mScarlet3 | Homologous recombination of aDM019; repair template: pDM154 (oDM528-531) | 4(A, B), S4(A-D), S5C |
| aDM084 | pyrG89; nkuA::argB?, scΔ?; ssdA::mScarlet3, GFP::tubA | Cross between aDM019 and XX872 | 3(A-C), S3(A-C) |
| aDM020 | pyrG89; fabM::mScarlet3; pyroA4, nkuA::argB, scΔ | Previous work (Modaffari et al., 2024) | S2(A-C, L) |
| aDM083 | pyrG89; fabM::mScarlet3; pyroA4, nkuA::argB, scΔ; ssdA::mNeonGreen | CRISPR edit of aDM020 using pDM044 gRNA; repair template: pDM155 (oDM664/665) | 2(A, B), S2(D, E) |
| aDM087 | pyrG89; fabM::mScarlet3; pyroA4, nkuA::argB, scΔ; ssdAΔ::NAT | CRISPR edit of aDM059 using pDM060 gRNA; repair template: pDM042 (oDM227/228) | S2(J-L) |

**Table S2.** Plasmids used in this study.

| Identifier | Description | Origin |
|---|---|---|
| CRISPR-Cas9 vector plasmid | | |
| pDM026 (Addgene #216808) | Vector for *Aspergillus* CRISPR-Cas9 genetic engineering with Golden Gate cloning drop-out cassette for spacer insertion and *pyrG* selectable marker. | Modaffari et al, 2024 |
| pDM068 (Addgene #216811) | Vector for Aspergillus CRISPR-Cas9 genetic engineering with Golden Gate cloning drop-out cassette for spacer insertion and *NAT* selectable marker. | Modaffari et al., 2024 |
| CRISPR-Cas9 sgRNA-carrying plasmids | | |
| pDM044 | sgRNA plasmid to target *ssdA* (AN1158); sgRNA1; spacer: AGACCCTTTACAGTGCGTAG; cloned into pDM026 (*pyrG* selection) | Modaffari et al., 2024 |
| pDM128 | sgRNA plasmid to target *ssdA* (AN1158); sgRNA5; spacer: AAGATAAGCCGAAGATCGTT; cloned into pDM026 (*pyrG* selection) | This study |
| pDM060 | sgRNA plasmid to target *fabM* (AN4000); sgRNA2; spacer: AGACAGAATTCCGACTTGCT; cloned into pDM026 (*pyrG* selection) | Modaffari et al., 2024 |
| pDM165 | sgRNA plasmid to target *ssdA* (AN1158); sgRNA1; spacer: AGACCCTTTACAGTGCGTAG; cloned into pDM068 (*NAT* selection) | This study |
| pDM137 | sgRNA plasmid to target *mobB* (AN1370); sgRNA2; spacer: ATAGTAAAAGGAGAGAAAGT; cloned into pDM026 (*pyrG* selection) | This study |
| Integration cassettes | | |
| pDM106 | *ssdA(AN1158)::pyrG* deletion cassette in pUC19 backbone; *A. fumigatus pyrG*; ~1kb homology arms | This study |
| pDM145 | *ssdA(AN1158)::NAT* deletion cassette in pUC19 backbone; *NAT* coding sequence is flanked by *A. fumigatus pyrG* promoter and terminator; ~1kb homology arms | This study |
| pDM112 | *pxdA(AN1156)::pyrG* deletion cassette in pUC19 backbone; *A. fumigatus pyrG*; ~1kb homology arms | This study |
| pDM154 | *dipA(AN10946)::pyrG* deletion cassette in pUC19 backbone; *A. fumigatus pyrG*; ~1kb homology arms | This study |
| pDM111 | Full *cotA-as2* (M281) gene with ~1kb homology arms; used for pDM116 construction | This study |
| pDM116 | *cotA-as2* (M281) end-in recombination plasmid; the construct contains the coding sequence of $CotA^{M281A}$ minus the start codon, followed by ~1kb sequence 3′ to stop codon and *A. fumigatus pyrG;* pUC19 backbone | This study |
| pDM159 | *ssdA-NtermΔ(2-429Δ)::msScarlet3* with ~1 kb homology 5′ to start codon and 3′ to stop codon; for integration into s*sdA::pyrG A. nidulans;* pUC19 backbone | This study |
| pDM160 | *ssdA-6A(S26A,S126A,S183A,S255A,S300A,S339A)::mScarlet3* with ~1 kb homology 5′ to start codon and 3′ to stop codon; for integration into *ssdA::pyrG A. nidulans*; pUC19 backbone | This study |
| pDM161 | *ssdA(S26D,S126D,S183D,S255D,S300D,S339D)::mScarlet3* with ~1 kb homology 5′ to start codon and 3′ to stop codon; for integration into *ssdA::pyrG A. nidulans*; pUC19 backbone | This study |
| Fluorescent protein cassettes | | |
| pDM042 (Addgene #216812) | *A. nidulans* codon-adjusted linker-mScarlet3 fluorescent protein, includes linker for C-terminal tagging; linker sequence: QAAALELVDP; pUC19 backbone | Modaffari et al., 2024 |
| pDM052 (Addgene #216814) | *A. nidulans* codon-adjusted linker-mNeonGreen fluorescent protein, includes linker for C-terminal tagging; linker sequence: QAAALELVDP; pUC19 backbone | Modaffari et al., 2024 |
| pDM155 | *A. nidulans* codon-adjusted 3xmNeonGreen. Construct: FLAG-(SGGS)x2-XTEN16-(GGGGS)x3-mNeonGreen-XTEN16-mNeonGreen-XTEN16-mNeonGreen-(GGGGS)x3-XTEN16-(SGGS)x2-FLAG; pUC19 backbone | This Study |

**Table S3.** Oligonucleotides and synthetic gene fragments used in this study.

| Identifier | Sequence | Description | Purpose |
|---|---|---|---|
| Oligonucleotides | | | |
| oDM045 | AGTGGGGATGCCTCAATTGTG | Amplify pyrG cassette. Reverse. | Plasmid construction (pDM112, pyrG cassette) |
| oDM133 | ctagacacctgcagcgggacAGACCCTTTA CAGTGCGTAGgtttcgaggcaggtgcttcc | To anneal with complementary oligo and Golden Gate using PaqCI into CRISPR plasmid. Spacer: ssdA_AN1158_gRNA1 (AGACCCTTTACAGTGCGTAG). Forward. | gRNA spacer (ssdA_AN1158_gRNA1, pDM044/pDM165) |
| oDM134 | ggaagcacctgcctcgaaacCTACGCACTG TAAAGGGTCTgtcccgctgcaggtgtctag | To anneal with complementary oligo and Golden Gate using PaqCI into CRISPR plasmid. Spacer: ssdA_AN1158_gRNA1 (AGACCCTTTACAGTGCGTAG). Reverse. | gRNA spacer (ssdA_AN1158_gRNA1, pDM044/pDM165) |
| oDM154 | TTGAGCAAACTCTGATCGCCTG | To sequence-verify correct insertion of protospacer in sgRNA plasmids. Binds A. fumigatus U3 terminator. Reverse. | gRNA plasmid verification (protospacer sequencing) |
| oDM185 | GAGCCCTCCGTAAGTTACGTTCGATTCTTA AACGCATTCTTCACTAACAATACATAATAT AGTTGCCTTACTATCCGCTCAGTCAACCCC TACGCACTGcagGCTGCTGCTctcgagctc | To amplify mScarlet3 from pDM042 for C terminal tagging of ssdA (AN1158). Includes ~100bp homology arm overhang. Forward. | Repair template amplification (ssdA::mScarlet3 from pDM042) for aDM094 |
| oDM186 | AGCATCATCACCAGATGGGCCCAAAGTCGG TGAAGCATGTAATATTCTTTGATGTAAAAC GTGTCGATTATGAGAGGTTAAGGATATAAA AGACCCTttaggaaccaccagaaccaccgg | To amplify mScarlet3 from pDM042 for C terminal tagging of ssdA (AN1158). Includes ~100bp homology arm overhang. Reverse. | Repair template amplification (ssdA::mScarlet3 from pDM042) for aDM094 |
| oDM193 | AGCATGCCACCCTCTCCTACCC | Genotyping and sequencing of ssdA (AN1158) tagged strains. Forward. | Genotyping/sequencing (ssdA C-term tagged strains; aDM019, aDM059, aDM094) |
| oDM194 | CACCACGTCGGACATGCTCACC | Genotyping and sequencing of ssdA (AN1158) tagged strains. Reverse. | Genotyping/sequencing (ssdA C-term tagged strains; aDM019, aDM059, aDM094) |
| oDM244 | CCCTAAGATCCAGGCCACTCAGC | Genotyping and sequencing of fabM (AN4000) tagged strains. Forward. | Genotyping/sequencing (fabM tagged strains; aDM087) |
| oDM245 | GTCTCAGCTACGAATGGTCGCAAG | Genotyping and sequencing of fabM (AN4000) tagged strains. Reverse. | Genotyping/sequencing (fabM tagged strains; aDM087) |
| oDM324 | AGGATCCCGCATACCTATACCTCCTGgccG AGTTCTTACCTGGAGGTGATTTGATGACCA | Amplify cotA and introduce M281A (as2) mutation. Used for construction of pDM111. Forward. | Plasmid construction (pDM111, cotA M281A mutagenesis) |
| oDM325 | TGGTCATCAAATCACCTCCAGGTAAGAACT CggcCAGGAGGTATAGGTATGCGGGATCCT | Amplify cotA and introduce M281A (as2) mutation. Used for construction of pDM111. Reverse. | Plasmid construction (pDM111, cotA M281A mutagenesis) |
| oDM459 | GAGCTAGAGCCCCCAGGACTAC | To amplify cotA (AN5529). Forward. | Genotyping aDM062 for correct integration of pDM116 |
| oDM202 | GCCAGCAACGCGGCCTTTTTAC | To amplify cotA (AN5529). Reverse. | Genotyping aDM062 for correct integration of pDM116 |
| oDM408 | TCCTCTAGAGTCGACCTGCAG | To amplify pUC19 for Gibson assemblies. Forward. | Plasmid construction (pUC19 backbone for pDM106/145/159/160/161) |
| oDM409 | TCCCCGGGTACCGAGCTC | To amplify pUC19 for Gibson assemblies. Reverse. | Plasmid construction (pUC19 backbone for pDM106/145/159/160/161) |
| oDM439 | TGAATTCGAGCTCGGTACCCGGGGATTGGG CCGTTCCTTCAAGTT | To amplify region upstream ssdA (AN1158) CDS for homology arm of repair template. Adds 25bp homology to pUC19 amplified with oDM408-409 for Gibson assembly. Forward. | Plasmid construction (pDM106, ssdA upstream homology arm) |
| oDM440 | CTTTGCAATCGGAGAGGGAG | to amplify ssdA upstream homology arm; also used in construction of ssdA N-terminal mutants plasmid. Reverse. | Plasmid construction (pDM106, ssdA upstream homology arm) |
| oDM441 | CTCCCTCTCCGATTGCAAAGACTAGGTAAT ATGACATGATTACGAATTCG | To amplify pyrG cassette from pDM023 with 20 bp homology to region upstream of ssdA CDS for Gibson assembly. Forward. | Plasmid construction (pDM106, pyrG cassette amplification) |
| oDM442 | TATGAACTAAAGTTTGGACTAGTGGGGATG CCTCAATTGT | To amplify pyrG cassette from pDM023 with 20 bp homology to region downstream of ssdA's last exon for Gibson assembly. Reverse. | Plasmid construction (pDM106, pyrG cassette amplification) |
| oDM443 | AGTCCAAACTTTAGTTCATATCCACAA | to amplify ssdA downstream homology arm; also used in construction of ssdA N-terminal mutants plasmid. Forward. | Plasmid construction (pDM106, ssdA downstream homology arm) |
| oDM444 | ATGCCTGCAGGTCGACTCTAGAGGAATCCC GGGACTCAGAGAACA | To amplify region downstream ssdA(AN1158)'s last exon for homology arm of repair template. Adds 25bp homology to pUC19 amplified with oDM408-409 for Gibson assembly. Reverse. | Plasmid construction (pDM106, ssdA downstream homology arm) |

| | | | |
|---|---|---|---|
| oDM445 | TGAATTCGAGCTCGGTACCCGGGGAATCGA GTGGACCAAGGCAAG | Binds ~1kb upstream of CotA start codon. For Gibson assembly of pDM111. Forward. | Plasmid construction (pDM111/116, cotA upstream region) |
| oDM450 | ATGCCTGCAGGTCGACTCTAGAGGATCGAA TGCCGTCTCTGTCTG | Binds ~1kb downstream of CotA stop codon. For Gibson assembly of pDM111. Reverse. | Plasmid construction (pDM111/116, cotA downstream region) |
| oDM461 | TTTCCTGCTTGCAAGTTCGC | Genotying ssdA deletion and mutants, Binds ~1.7kb upstream of ssdA start codon. Forward. | Genotyping/sequencing (ssdA deletion/mutants; aDM039, aDM059, aDM078, aDM080, aDM086, aDM089) |
| oDM462 | AGCACTGCATGAAGACCTCC | Genotying ssdA deletion. Binds ~1.8kb downstram of ssdA stop codon. Reverse. | Genotyping/sequencing (ssdA deletion/mutants; aDM039, aDM059, aDM078, aDM080, aDM086, aDM089) |
| oDM474 | TGAATTCGAGCTCGGTACCCGGGGATCAGG CCCGACTACGATGAG | Amplify pxdA (AN1156) upstream homology arm for Gibson assembly. Forward. | Plasmid construction (pDM112, pxdA upstream homology arm) |
| oDM475 | CGAATTCGTAATCATGTCATATTACCTAGT CACTGGTCGCTTGAGGGAGGT | Amplify pxdA (AN1156) upstream homology arm for Gibson assembly. Reverse. | Plasmid construction (pDM112, pxdA upstream homology arm) |
| oDM476 | ACTAGGTAATATGACATGATTACGAATTCG | Amplify pyrG cassette. Forward. | Plasmid construction (pDM112, pyrG cassette) |
| oDM477 | CTAGCACAATTGAGGCATCCCCACTGATTC TGATTTCACAAACTACCTGTCTCTG | Amplify pxdA (AN1156) downstream homology arm for Gibson assembly. Forward. | Plasmid construction (pDM112, pxdA downstream homology arm) |
| oDM478 | ATGCCTGCAGGTCGACTCTAGAGGACTTGG TCCCCAGCTTACCAG | Amplify pxdA (AN1156) downstream homology arm for Gibson assembly. Reverse. | Plasmid construction (pDM112, pxdA downstream homology arm) |
| oDM520 | ACAGCCCCAGAGCTATCAGA | To amplify pxdA (AN1156) region outside of homology arms for genotyping and sequencing deletion. Forward. | Genotyping/sequencing (pxdA deletion; aDM049) |
| oDM521 | TTGAAGCCCCTCGATCCCTA | To amplify pxdA (AN1156) region outside of homology arms for genotyping and sequencing deletion. Reverse. | Genotyping/sequencing (pxdA deletion; aDM049) |
| oDM527 | CGTGGAGTTACCAGTGATTGACC | Amplify pyrG cassette. Forward. | Plasmid construction (pDM154, pyrG cassette) |
| oDM528 | TTGGCCTATACGAGGGCAAC | Amplify dipA (AN10946) upstream homology arm for Gibson assembly. Forward. | Plasmid construction (pDM154, dipA upstream homology arm) |
| oDM529 | GTAGAAGATAAAACATTGGTCAATCACTGG TAACTCCACGTGGTAGACGAGGTGCAACAG | Amplify dipA (AN10946) upstream homology arm for Gibson assembly. Reverse. | Plasmid construction (pDM154, dipA upstream homology arm) |
| oDM530 | GCAAGGCGGCCGCAGCTAGCACAATTGAGG CATCCCCACTAACCAATTCCACGGTTCCCA | Amplify dipA (AN10946) downstream homology arm for Gibson assembly. Forward. | Plasmid construction (pDM154, dipA downstream homology arm) |
| oDM531 | GATTTTAGGCGAATGGGCGG | Amplify dipA (AN10946) downstream homology arm for Gibson assembly. Reverse. | Plasmid construction (pDM154, dipA downstream homology arm) |
| oDM536 | CAGAAGTGAAGACGCTCAGAATGGTGTTTA TGGTCAG | Amplify cotA CDS for cotA-as2 plasmid Gibson assembly. Forward, | Plasmid construction (pDM116, cotA CDS amplification) |
| oDM538 | CTACCCAGAGCGCTTCTCTG | To linearise cotAas2 plasmid pDM116 from end-in recombination. Forward. | Repair template linearisation (pDM116 for cotA-as2; aDM062) |
| oDM539 | CGTTGAAGTTCTTCGTTCTCG | To linearise cotAas2 plasmid pDM116 from end-in recombination. Reverse. | Repair template linearisation (pDM116 for cotA-as2; aDM062) |
| oDM546 | CTAGACACCTGCAGCGGGACAAGATAAGCC GAAGATCGTTGTTTCGAGGCAGGTGCTTCC | To anneal with complementary oligo and Golden Gate using PaqCI into CRISPR plasmid. Spacer: ssdA_AN1158_gRNA5 (AAGATAAGCCGAAGATCGTT). Forward. | gRNA spacer (ssdA_AN1158_gRNA5, pDM128) |
| oDM547 | GGAAGCACCTGCCTCGAAACAACGATCTTC GGCTTATCTTGTCCCGCTGCAGGTGTCTAG | To anneal with complementary oligo and Golden Gate using PaqCI into CRISPR plasmid. Spacer: ssdA_AN1158_gRNA5 (AAGATAAGCCGAAGATCGTT). Reverse. | gRNA spacer (ssdA_AN1158_gRNA5, pDM128) |
| oDM557 | TATCCACGATATGGGCACGC | Binds ~1.1kb upstream of dipA (AN10946) start codon; for genotyping and sequencing. Forward. | Genotyping/sequencing (dipA deletion; aDM056) |
| oDM558 | CCCGCTGCTATTTGGAGACT | Binds ~1.6kb downstreav of dipA (AN10946) stop codon; for genotyping and sequencing. Reverse. | Genotyping/sequencing (dipA deletion; aDM056) |
| oDM583 | CTAGACACCTGCAGCGGGACATAGTAAAAG GAGAGAAAGTGTTTCGAGGCAGGTGCTTCC | To anneal with complementary oligo and Golden Gate using PaqCI into CRISPR plasmid. Spacer: mobB_AN1370_gRNA2 (ATAGTAAAAGGAGAGAAAGT). Forward. | gRNA spacer (mobB_AN1370_gRNA2, pDM137) |

| | | | |
|---|---|---|---|
| oDM584 | GGAAGCACCTGCCTCGAAACACTTTCTCTC<br>CTTTTACTATGTCCCGCTGCAGGTGTCTAG | To anneal with complementary oligo and Golden Gate using PaqCI into CRISPR plasmid. Spacer: mobB_AN1370_gRNA2 (ATAGTAAAAGGAGAGAAAGT). Reverse. | gRNA spacer (mobB_AN1370_gRNA2, pDM137) |
| oDM587 | ATCCACCGCCCATTCGCCTAAAATCGGAGA<br>GGCGGTTTGCGTATTGGG | Amplify pUC19 backbone with dipA downstream homology arm homology for Gibson assembly. Forward. | Plasmid construction (pDM154, pUC19 backbone) |
| oDM588 | CTCCTGTTGCCCTCGTATAGGCCAAGCGCA<br>GCCTGAATGGCGAATGG | Amplify pUC19 backbone with dipA downstream homology arm homology for Gibson assembly. Reverse. | Plasmid construction (pDM154, pUC19 backbone) |
| oDM616 | GGTGGTCACTGGGGCCAGCAGAGCTCTAGC<br>CAAGCTGCTGCTTTAGAATTAGTTGATCC | To amplify mNeonGreen from pDM052 for tagging mobB (AN1370) using pDM137. Forward. | Repair template amplification (mobB::mNeonGreen from pDM052) to construct aDM070 |
| oDM617 | AAGGAGAGAAAGTGGtCGATATACTCTAGC<br>TTActtgtagagctcgtccataccca | To amplify mNeonGreen from pDM052 for tagging mobB (AN1370) using pDM137. Introduces C->A PAM mutation in 3' UTR. Reverse. | Repair template amplification (mobB::mNeonGreen from pDM052) to construct aDM070 |
| oDM635 | CGGCAACAAGATAAGCCGAAGATCGTTGCC<br>TTCGCTCCTACCGACAAGCGTGTGCCTTTG | To anneal with oDM636 and use as a repair template for SsdA-WAKA (W652A, K654A) with pDM128 gRNA plasmid. Forward. | Repair template (ssdA W652A/K654A annealed oligos) to construct aDM073 |
| oDM636 | CAAAGGCACACGCTTGTCGGTAGGAGCGAA<br>GGCAACGATCTTCGGCTTATCTTGTTGCCG | To anneal with oDM636 and use as a repair template for SsdA-WAKA (W652A, K654A) with pDM128 gRNA plasmid. reverse. | Repair template (ssdA W652A/K654A annealed oligos) to construct aDM073 |
| oDM637 | ACTCAGAACGCTCCCTCTCCGATTGCAAAG<br>atgACTCTGTTCACTCCCTATCTTCCCC | To amplify SsdA-mScarlet3 from aDM019 for Gibson asembly of N-term deletion cassette. Forward. | Plasmid construction (pDM159, SsdA-mScarlet3 N-term deletion) |
| oDM638 | GATTTGTGGATATGAACTAAAGTTTGGACT<br>CCTAAAAGGGACTATTTGTCGCCATTCGG | To amplify SsdA-mScarlet3 from aDM019 for Gibson asembly of N-term deletion cassette. Reverse. | Plasmid construction (pDM159, SsdA-mScarlet3 N-term deletion) |
| oDM645 | ATGAACCAGTCAATCGGCTC | To amplify SsdA-mScarlet3 from aDM019 for Gibson asembly of phosphomutants. Forward. | Plasmid construction (pDM160/161, SsdA phosphomutants) |
| oDM646 | GCCACACTAGGATTTGTGGATATG | To amplify SsdA-mScarlet3 from aDM019 for Gibson asembly of phoshpomutants. Reverse. | Plasmid construction (pDM160/161, SsdA phosphomutants) |
| oDM647 | GAAATGGAGCCAATGCAGCC | To amplify around C term of MobB (AN1370) for genotyping and sequencing. Forward. | Genotyping/sequencing (mobB C-term tagged; aDM070) |
| oDM648 | TCGCATCATCTGCCTATCCG | To amplify around C term of MobB (AN1370) for genotyping and sequencing. Reverse. | Genotyping/sequencing (mobB C-term tagged; aDM070) |
| oDM649 | GAAGCCCCTATACGCTGGTC | To amplify around ssdA-WAKA mutant residues for genotyping and sequencing. Forward. | Genotyping/sequencing (ssdA-WAKA; aDM073) |
| oDM651 | CCCGACGGGAAGACAAATCA | To amplify around ssdA-WAKA mutant esidues for genotyping and sequencing. Reverse. | Genotyping/sequencing (ssdA-WAKA; aDM073) |
| oDM664 | ACTATCCGCTCAGTCAACCCCTACGCACTG<br>tccggtggttctagcggtg | To amplify 1xmNeonGreen from pDM155 for tagging of SsdA (AN1158) using gRNA1, 30bp homology arm. Forward. | Repair template amplification (ssdA::mNeonGreen from pDM155) to construct aDM083 and aDM088 |
| oDM665 | TGAGAGGGTAAGGATATAAAAGACCCCTTTA<br>cttatacagctcgtccattcccatg | To amplify 1xmNeonGreen from pDM155 for tagging of SsdA (AN1158) using gRNA1, 30bp homology arm. Reverse. | Repair template amplification (ssdA::mNeonGreen from pDM155) to construct aDM083 and aDM088 |
| oDM535 | ACCATTCTGAGCGTCTTCACTTCTGAGTTC<br>GGCATGG | Amplify pUC19 backbone for cotA-as2 plasmid Gibson assembly. Reverse. | Plasmid construction (pDM116, pUC19 backbone) |
| Synthetic gene fragments | | | |

| | | | |
|---|---|---|---|
| gDM018 | gacaaatccggtggttctagcggtggttctatggtgtccaaaggagaagaggacaatatggcctctctccccgctactca<br>cgagctccacatctttggaagcatcaatggtgttgacttcgatatggtcggtcaaggtacaggtaatcccaatgacggat<br>acgaagaactcaatctcaagtctacaaagggtgatctccagttctctccgtggatccttgtcccgcatatcggatatggc<br>ttccaccagtacctcccctatccggatggcatgtctccctttcaagctgcgatggtcgacggctctggataccaggttca<br>ccgtaccatgcaattcgaggacggtgcttctctgactgtcaactaccgttatacatacgagggctcccatattaagggcg<br>aggcccaggtgaaaggtacgggatttcctgctgatggacccgttatgactaactccctcaccgccgctgattggtgtcgc<br>tccaagaaaacctacccgaatgacaagaccattatttccaccttcaagtggtcctacaccactggaaacggtaaacgtta<br>ccgcagcaccgctcgcaccacctatacattcgccaaacctatggctgcgaactaccttaagaaccaacctatgtacgtct<br>ttcgcaagaccgaactgaaacactctaagacggagctcaattttaaggagtggcagaaggcgttcactgacgtcatggga<br>atggacgagctgtataagtccggttccgagactcctggtacctctgaaagcgctactcccgagtctgtctctaagggaga<br>ggaggacaacatggctagccttcctgcgacccatgagctgcacattttcggctccattaacggagtcgatttcgacatgg<br>tcggacagggtaccggaaaccctaacgacggttacgaggaacttaatctgaagtccaccaaaggcgaccttcagtttagc<br>ccctggatcctcgttcctcacattggctacggtttccatcaatacctgccttatcccgacggtatgagcccgtttcaagc<br>cgctatggtggatggttccggctaccaagttcatcgcacaatgcagtttgaggatggtgcgtctctcaccgtgaattacc<br>gctatacctatgagggttcccacatcaaaggtgaagcgcaagtcaagggcaccggttttcctgctgatggacctgtcatg<br>accaacagccttactgccgccgactggtgtcgtagcaaaaagacctatcccaacgataagacgatcatctccacgtttaa<br>gtggtcttacacaaccggcaacggcaagcgttatcgttctactgcccgcactacatataccttcgctaaacccatggcgg<br>ctaactacctgaagaaccagccgatgtatgtcttccgtaaaacggaactcaagcacagcaagacagagcttaacttcaaa<br>gagtggcagaaagccttcacagatgtgatgggtatggacgaactgtacaagtctggtagcgaaacccccggtacttccga<br>atctgctaccccctgagtccgttagcaagggcgaagaagataacatggccagccttcccgccactcatgaactccatatct<br>tcggttctatcaacggcgtcgactttgacatggtgggccaaggcactggtaacccgaacgatggttatgaggagctgaac | *A. nidulans* codon-adjusted 3x mNeonGreen; XTEN16 linker in between mNeonGreen copies; contains homology for Gibson assembly into pDM097 | Construction of pDM155 |

| | | | |
|---|---|---|---|
| | ctcaagagcactaagggtgacctccaattctccccttggattctggtgccccacattggttacggctttcatcagtatct<br>tccctaccccgatggtatgtccccttttcaggctgccatggttgacggttctggttatcaggtgcatcgtactatgcagt<br>tcgaagatggcgcctccctgaccgttaactatcgttacacctacgaaggttctcatatcaagggtgaggctcaggtcaaa<br>ggaactggtttcccggcggatggtcctgtgatgacaaactctcttacagctgccgattggtgccgttctaagaagactta<br>ccccaacgacaaaaccatcattagcacattcaagtggagctacacgaccggtaatggtaaacgctaccgctctacggccc<br>gtactacctacacttttgcgaagcctatggccgccaattacctcaaaaatcagcccatgtacgttttccgcaagactgag<br>ctcaaacattccaaaaccgagctcaacttcaaggaatggcaaaaggccttcaccgacgttatgggcatggatgagctcta<br>caagggtggtggtggttctggtggtggcggatct | | |
| gDM019 | GGTATCTCACACTCAGAACGCTCCCTCTCCGATTGCAAAGATGGATCAACAGAATCAGTCCGGGGTCCCCGGACCTGCTG<br>GACGAAAGCTTCATATTGCCCATAGACGGAGCCCAGCTGAGCTTACCCCTCTCATGAGTATGTTCCGCTCACAATTTATT<br>GATCTGCTCGCCTTCAAGTAGTTTGAGATGCTAACTGTCAGATCTACAGTGGAACAACTGGCGATCCAGCAGCAAATTGA<br>GTTGCTTCAACAGCAACAACAGCAAATTGCAGCTACACACCAGCAGTATATCAATATGGGTTTATTGCAACCGCAGCAAA<br>TCGGCCAGGCTTCTGGTTTCACGCCGTCTCTACAAGGTGGCGGTGCGCCCATCGGCGGCGTTTCTCCACAACAACTCGGT<br>GCCTTTCAGTTTCCTCAGGTTGGCCAGCAGCAGCAGCTAGGCGTTCCGGGGAACACTTCGAATCAGCATTCGCATCGCCG<br>TAATCAAGCTGCTCTCCCAGGCTTCGGAATGGGCGGTCCTCCCCCAGCTCCATCTTCTGGAGCTTCGGGTTATTCGGACT<br>ACAGCCAACAGCAACAGGGTAACTTCCAAAGGAACGAGAACGGGGCTCATGGCCGTGGCCGTGGGGGCCCTTCGGGCGGT<br>GGTCACCAGCGACGTCATGCTCTTGCTCTCCCTGAAGCGAAGAAGGCTGCGGAGCTCGCTCAGCAAAAGCGGGCGGCATC<br>TGGGTTCCAATTCCCTGCCCCCACCGGTGACTCGTCTGAGGCCGCACCCGCATCCGACGACAAACCAAGCACCCCGAATC<br>CAACACAGTCGTCTCAAGGATTGGGTCTTCAGCGTGCCGGCAACATCAGAGCCAGTGCTCATGGCCGCAGTCAGGCAATG<br>GCTGTGGGTGGTAACCGCGGCTCTTTGTCTGGTCGTGGTCCTGGTGGTTTTCAATTCCCTGCGAATGATGGTTCCTCTGG<br>CCCTGAGAACCAACGCCGAGGCAGCCAAGCGTCCCACGCCCGAACCCCAGCTAGGAACTTTGACGGCAACTGGCGCCAGC<br>CCAACAACCAATCTCAAGGCCAGGACCAATCTAAAGGTCTCGGACAGCAGAACTCAGGATTTCAGCCCGGACACCGTGCA<br>CGTGCGGCTATGAACCAGTCAATCGGCTCTATCGGTTCCTTCCAATACC | N-terminal region of *ssdA* with nonphosphorylatable mutations at predicted Cbk1/NDR sites (S26A, S126A, S183A, S255A, S300A, S339A) | Construction of pDM160 |
| gDM020 | GGTATCTCACACTCAGAACGCTCCCTCTCCGATTGCAAAGATGGATCAACAGAATCAGTCCGGGGTCCCCGGACCTGCTG<br>GACGAAAGCTTCATATTGCCCATAGACGGAGCCCAGACGAGCTTACCCCTCTCATGAGTATGTTCCGCTCACAATTTATT<br>GATCTGCTCGCCTTCAAGTAGTTTGAGATGCTAACTGTCAGATCTACAGTGGAACAACTGGCGATCCAGCAGCAAATTGA<br>GTTGCTTCAACAGCAACAACAGCAAATTGCAGCTACACACCAGCAGTATATCAATATGGGTTTATTGCAACCGCAGCAAA<br>TCGGCCAGGCTTCTGGTTTCACGCCGTCTCTACAAGGTGGCGGTGCGCCCATCGGCGGCGTTTCTCCACAACAACTCGGT<br>GCCTTTCAGTTTCCTCAGGTTGGCCAGCAGCAGCAGCTAGGCGTTCCGGGGAACACTTCGAATCAGCATTCGCATCGCCG<br>TAATCAAGACGCTCTCCCAGGCTTCGGAATGGGCGGTCCTCCCCCAGCTCCATCTTCTGGAGCTTCGGGTTATTCGGACT<br>ACAGCCAACAGCAACAGGGTAACTTCCAAAGGAACGAGAACGGGGCTCATGGCCGTGGCCGTGGGGGCCCTTCGGGCGGT<br>GGTCACCAGCGACGTCATGACCTTGCTCTCCCTGAAGCGAAGAAGGCTGCGGAGCTCGCTCAGCAAAAGCGGGCGGCATC<br>TGGGTTCCAATTCCCTGCCCCCACCGGTGACTCGTCTGAGGCCGCACCCGCATCCGACGACAAACCAAGCACCCCGAATC<br>CAACACAGTCGTCTCAAGGATTGGGTCTTCAGCGTGCCGGCAACATCAGAGCCAGTGCTCATGGCCGCAGTCAGGACATG<br>GCTGTGGGTGGTAACCGCGGCTCTTTGTCTGGTCGTGGTCCTGGTGGTTTTCAATTCCCTGCGAATGATGGTTCCTCTGG<br>CCCTGAGAACCAACGCCGAGGCAGCCAAGCGTCCCACGCCCGAACCCCAGACAGGAACTTTGACGGCAACTGGCGCCAGC<br>CCAACAACCAATCTCAAGGCCAGGACCAATCTAAAGGTCTCGGACAGCAGAACTCAGGATTTCAGCCCGGACACCGTGCA<br>CGTGCGGACATGAACCAGTCAATCGGCTCTATCGGTTCCTTCCAATACC | N-terminal region of *ssdA* with phosphomimetic mutations at predicted Cbk1/NDR sites (S26D, S126D, S183D, S255D, S300D, S339D) | Construction of pDM161 |
